# Cooperative progression of colitis and leukemia modulated by clonal hematopoiesis via PTX3/IL-1β pro-inflammatory signaling

**DOI:** 10.1101/2023.08.29.555330

**Authors:** Hang He, Yuchen Wen, Hanzhi Yu, Jingjing Liu, Qingran Huo, Wenyan Jin, Zhiqin Wang, Guohui Du, Jun Du, Huaquan Wang, Zhigang Zhao, Zhigang Cai

## Abstract

Clonal hematopoiesis (CH) is considered an important risk factor for all-cause mortality and the development of multiple chronic diseases including hematological neoplasms, cardiovascular diseases, and potentially a range of autoimmune or immune-deficiency diseases. Mutations in *TET2* are one of the first identified, most important, and prevalent genetic drivers of CH. However, cooperative factors and mechanisms underlying *TET2*-deficiency related CH (TedCH) remain largely unknown. Recently, it has been suggested that certain diseases occurred before TedCH and promote TedCH trajectory on the contrary, indicating that diseases in non-hematopoietic organs may act as environmental non-genetic drivers of CH. To clarify the relationships between immune-dysfunctional diseases and CH, here we tested the impact of various challenges on TedCH. We found that expedited TedCH depended on establishment of an inflammatory environment. Primary or chimeric *Tet2*-mutant mice spontaneously developed co-symptoms reminiscent of human chronic colitis and myeloid leukemia, which was exacerbated by feeding with DSS, an experimental inducer of ulcerative colitis. Single cell RNA-seq (scRNA-seq) analysis reveals in depth the damage of colon in the *Tet2*-mutant mice in physiological conditions or fed with DSS, along with increase of dysbacteriosis indicated by gut microbiome analysis. Results from colon scRNA-seq from both mouse and human highlight the important roles of PTX3/IL-1β pro-inflammatory signaling in promoting colitis or leukemia. Finally, TedCH trajectory and inflammation in colon and bone marrow were ameliorated by treatment of IL-1R1 inhibitor Anakinra. Our study suggests that PTX3/IL-1β signaling and clonal hematopoiesis cooperate and play important roles in gut-bone marrow axis and related diseases including colitis and leukemia.

**Highlights:**

1. Certain environmental factors, such as Dextran Sulfate Sodium (DSS), an experimental inducer of ulcerative colitis, promote TedCH
2. Colitis and leukemia are spontaneously and simultaneously developed in *Tet2*-defficient primary or chimeric mice, along with increased pathogenic gut microbiomes, indicating an aberrant gut-bone marrow axis in the mutant mice.
3. Single cell RNA-seq analysis reveals enhanced PTX3, a soluble pattern recognition molecule and IL-1β pro-inflammatory signaling in colitis and leukemia.
4. The *In vivo* function of the PTX3/IL-1β pro-inflammatory signaling in TedCH is indicated in human colitis and validated in experimental settings.

## Introduction

Clonal hematopoiesis (CH) is a phenomenon in which hematopoietic stem cells (HSCs) carry genetic mutations with advantageous growth potential, resulting in aberrant expansion of certain immature and mature hematopoietic cell populations over time (Genovese et al., 2014; Steensma et al., 2015). Similar phenomenon, somatic mutation-driven clonal expansion of certain cells, takes place in non-hematopoietic organs such as skin, esophagus, and brain (Jaiswal and Ebert, 2019). Recent studies suggest that CH is associated with an increased risk of developing hematological malignancies, including acute myeloid leukemia (AML) and myelodysplastic syndrome (MDS). Scoring CH is able to stratify high-risk CH-carriers (Weeks et al., 2022). Interestingly, CH is also associated with non-hematological diseases, including all-cause mortality, cardiovascular disease, chronic obstructive pulmonary disease, and gout (metabolic arthritis, an auto-immune disease in bone juncture) (Agrawal et al., 2022; Jaiswal and Libby, 2020; Jaiswal et al., 2017; Miller et al., 2022). Accordingly, World Health Organization recently recognizes that CH is a precursor lesion of myeloid neoplasms (Khoury et al., 2022). Therefore, characterizing drivers and consequences of CH will not only assist in investigating clonal expansion of somatic cells in non-hematopoietic organs but also help in understanding the etiology of numerous chronic diseases many years prior to their onset (Weeks et al., 2022).

Loss-of-function heterozygous mutations in the epigenetic regulator *TET2* (Ten-Eleven Translocation methyl-cytosine dioxygenase-2) were among the first discovered and one of the most prevalent and important drivers of CH. Unsurprisingly now, mutations in *TET2* are also the top prevalent single-gene mutations in MDS, an age-related hematological disease with abnormal hematopoietic stem and progenitor cells (HSPCs) (Delhommeau et al., 2009; Nazha et al., 2021). *Tet2*-deficient primary or chimeric mouse models display a competitive advantage over normal HSPCs, skewing myeloid differentiation, and a tendency to transform into full-blown leukemia when co-operative mutations and environmental factors involved in (e.g., *Flt3^ITD/+^* or *Nras^G12D^*) (Moran-Crusio et al., 2011) (Cai et al., 2020a; Chu et al., 2012). Loss of *TET2* results in increase of DNA methylation at various genomic sites, such as promoters, enhancers, and CpG islands (Rasmussen et al., 2015). Recently, we and others reported that *Tet2*-deficient HSPCs exhibit significantly increased expression of the inflammatory cytokine interleukin-6 (*IL-6*) and show stronger anti-apoptotic and self-renewal abilities after exposure to acute stimulus such as *IL-6* or lipopolysaccharides (LPS), leading to the development of an age-dependent chronic myelomonocytic leukemia (CMML)-like disease in the mutant mice over time (Cai et al., 2018; Zhang et al., 2015). Microbial signals are a major source of inflammatory inducer and interestingly, *Tet2* mutant mice with antibiotic treatment or raised in germ-free condition developed minimal CMML-like phenotypes (Meisel et al., 2018). In line with these studies, a most recent study based on *in vivo* models rather than generating chimeric *Tet2*-mutant models demonstrated that interleukin-1 (*IL-1α* and *IL-1β*), is able to act as an external factor in driving *Tet2*-deficiency related clonal hematopoiesis (TedCH) over age; genetic loss of their receptor *IL1-R1* mitigated *Tet2*-deficiency related abnormalities (Burns et al., 2022; Caiado et al., 2023). In summary, results from several independent studies strongly suggest that *Tet2*-deficient HSPCs manifest TedCH phenomenon and cooperate with intrinsic and extrinsic factors to develop stronger symptoms such as full-blown AML.

However, through mathematical analysis and experimental validation, a recent study of TedCH suggests that hypercholesterolemia-induced atherosclerosis took place prior to clonal hematopoiesis and may act as a driver of *Tet2*-deficient HSCs proliferation on the contrary. This study suggests that certain diseases or environmental conditions, for example, atherosclerosis drive the onset and trajectory of TedCH (Heyde et al., 2021). Atherosclerosis is a complicated and chronic disease where inflammation dysregulation explains in part (if not all) the etiology: mononuclear cells from the blood, including white blood cells, are recruited to the vessel wall in response to tissue damage (Libby et al., 2002). To further demonstrate that “diseases” were able to take prior to TedCH and drive TedCH, the authors presented an additional experimental model, a sleep fragmentation model, and demonstrated that sleep disturbance drives TedCH (Heyde et al., 2021). Nonetheless, clarifying the drivers and consequences of TedCH is the most critical challenge in the area. Controlling the trajectory of TedCH will also assist in developing strategies to mitigate CH-related diseases.

In previous studies, we have identified LPS, an inducer of innate-immunity related TLR4 signaling, and mutations in *Flt3*, *Ins2* or lncRNA *Morrbid* cooperates with *Tet2* deficiency to exacerbate or mitigate TedCH or leukemic phenotypes(Cai et al., 2020a; Cai et al., 2018; Cai et al., 2021; Cai et al., 2020b). To further explore the potential drivers of CH, we assessed the impact of another four different environmental factors on TedCH and confirmed that accelerated TedCH depends on the establishment of an inflammatory environment. Based on the results of single cell RNA-seq in the organs of colon (mouse and human both included) and microbe sequencing in gut, we propose that the PTX3/IL-1β signaling pathway, a tight complement/pro-inflammation cascade, plays important roles in gut-bone marrow axis which in turn regulates clonal hematopoiesis. We suggest that the PTX3/IL-1β signaling pathway should be appreciated to mitigate TedCH trajectory and related diseases including colitis and leukemia.

## Results

### DSS but not STZ, 5-FU, or additional irradiation promotes TedCH

Competitive bone marrow transplantation (cBMT) assays were developed for analyzing TedCH in the chimeric mice and four different treatments were chosen for testing environmental impacts on TedCH: DSS, STZ, 5-FU, and additional irradiation (**Figure 1A**). 5-FU and additional irradiation are well-known direct regulators of hematopoiesis while DSS induces colitis and systematic inflammation and STZ induces hypoglycemia. TedCH trajectory was monitored every 3 weeks by analyzing CD45.1, CD45.2 and CD45.1/CD45.2 chimerism in the peripheral blood (PB) of the chimeric mice. As shown in **Figure 1B** and **C**, expedited TedCH were observed upon DSS feeding compared to vehicle (Veh) feeding. Treatments with STZ or additional irradiation appear to have no effect on TedCH while 5-FU has significant inhibition on TedCH (**Figure 1D** to **F; Supplemental Figure 1A** to **C**). As DSS induces inflammation and colitis, we conclude that inflammation and colitis positively cooperate TedCH trajectory in the cBMT experimental setting.

**Figure 1.**
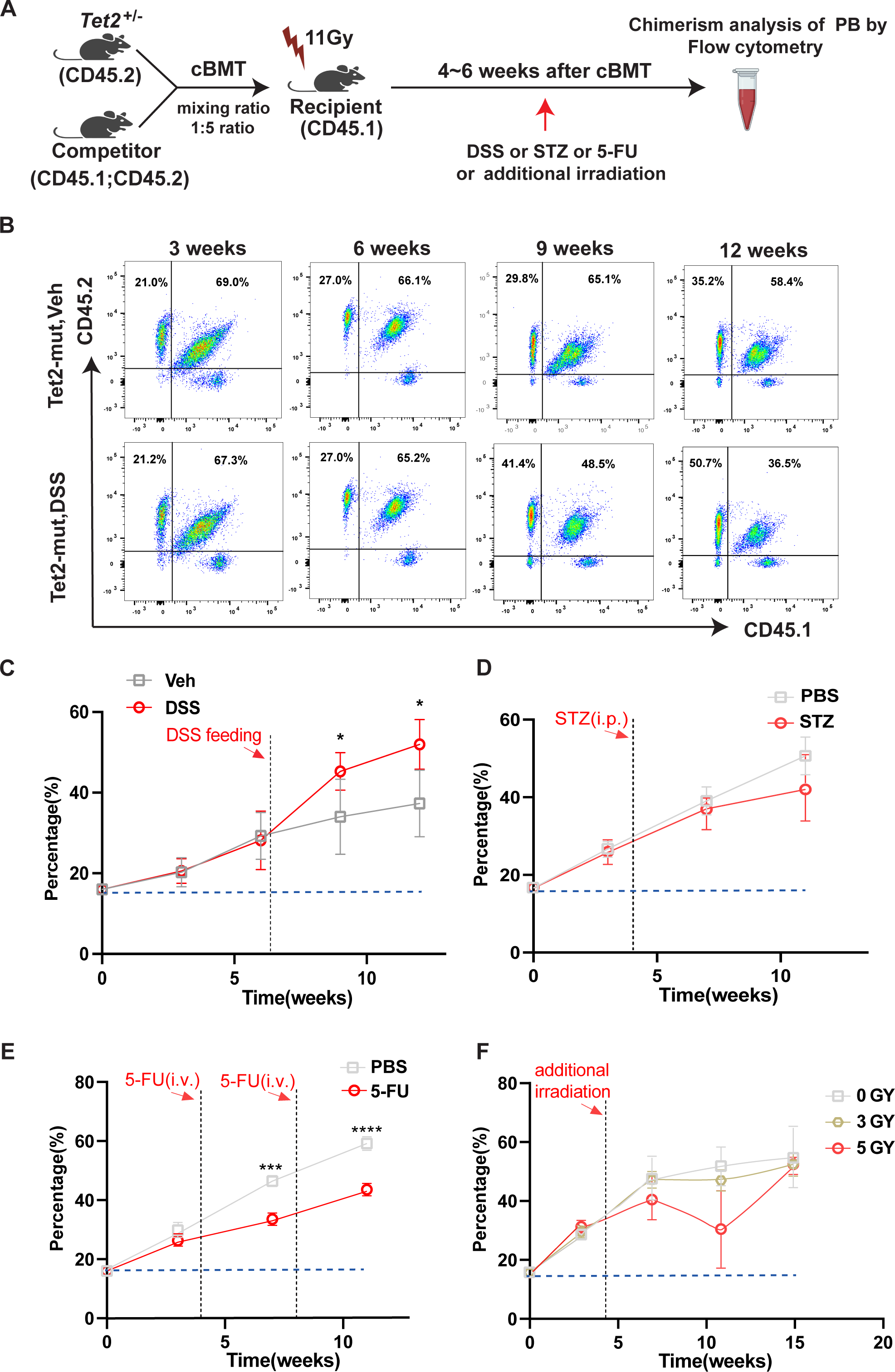
Assessing impacts of various treatments on TedCH trajectory. (A) Competitive bone marrow transplantation (cBMT) assays were used to search for cooperative environmental factors of TedCH. See Methods for detailed cBMT experimental procedures. Treatments were applied once TedCH is set up at the 3∼6 weeks post cBMT. Results of four different treatments (DSS, STZ, 5-FU and additional irradiation) were included in this study. As shown in the Figure 1A, DSS accelerated TedCH. Expedited TedCH was also observed when *Tet2^+/-^* HSPCs were mixed with genetically stable inflammatory HSPCs and the detail results will be reported in another study (Wen and Cai *et al*., manuscript in preparation, 2023). cBMT, competitive bone marrow transplantation; PB, peripheral blood. (B and C) Representative flow profiles of PB from the chimeric mice fed with DSS or normal water (vehicle, veh) (B) and the quantified trajectories of TedCH in the chimeric mice (C). (D-F) Quantified trajectories of TedCH in the chimeric mice treated with STZ (D), or 5-FU (E), or additional irradiation (F) as indicated. Data are shown as means ± SEMs. Number of biological repeats (animals): n = 5∼7. Experiments of DSS treatment were repeated twice. Experiments of STZ or 5-FU or additional irradiation treatment were performed once. *, p < 0.05; **, p < 0.01; ***, p < 0.001; ****, p < 0.0001. See also **Supplemental Figure 1** for representative flow profiles of PB from the chimeric mice with or without STZ, 5-FU and additional irradiation treatment.

### DSS exacerbates colitis burden and aberrant dysbacteriosis in *Tet2^+/-^* primary mice

Since DSS expedited TedCH in the chimeric mice, we then asked in detail what are the underlying biological and pathological processes and consequences during the DSS challenge. We chose *Tet2*-defficient primary mice for further study since they are more stable and readily monitored in a large cohort without irradiation involvement (irradiation itself may damage gut histology). A previous report suggests that acute DSS challenge exacerbates inflammatory bowel disease (IBD) in *Tet2*-defficient primary (Zhang et al., 2015). We turned to ask about the impact of chronic colitis by three cycles of low-dose DSS feeding (**Figure 2A**). Consistently, even low-dose of DSS also induced exacerbated colitis in *Tet2*-defficient primary mice (*Tet2^+/-^*_DSS vs. WT_DSS, **Figure 2B-E**). Upon DSS challenge, increased disease score (based on colon H&E staining) was observed and confirmed by expression of gut barrier markers *Muc2* and *Occludin* (**Figure 2F** and **G**; and **Supplemental Figure 2A**). Interestingly, *Tet2*-defficient primary mice appear to develop colitis spontaneously as shown by FITC-D staining in the serum and H&E staining in colon (*Tet2^+/-^*_Veh vs. WT_Veh, **Figure 2H**; and **Supplemental Figure 2A**).

**Figure 2.**
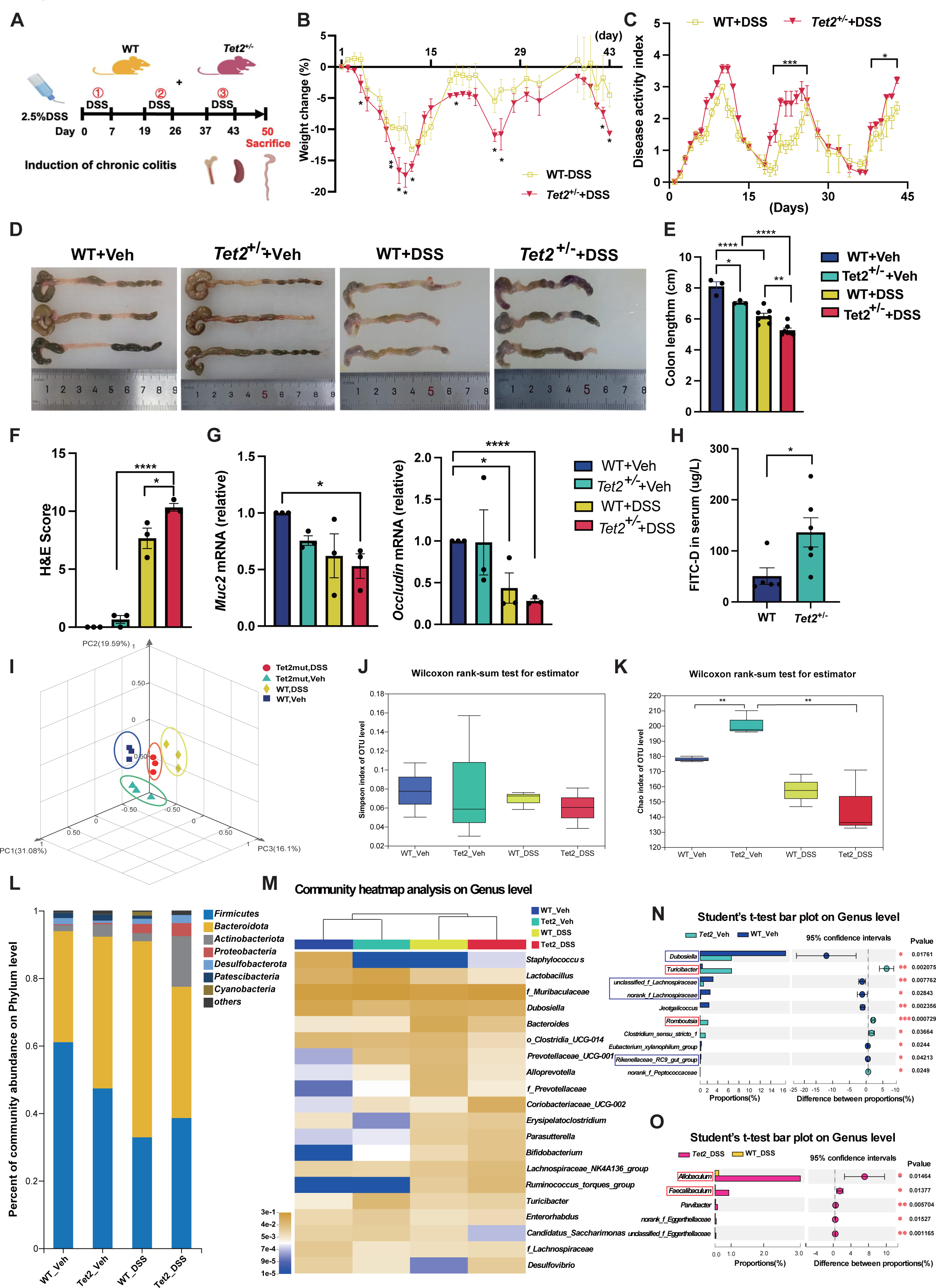
*Tet2*-deficient mice manifest exacerbated colitis and aberrant microbiota composition when fed with DSS. (A) Experimental scheme for induction of chronic colitis by DSS. Three cycles of DSS feeding were performed during the 50-day experimental procedure. Four groups of animals were included in the entire procedure: WT_Veh, *Tet2^+/-^*_Veh, WT_DSS, *Tet2^+/-^*_DSS. (B and C) Body weights and disease scores (disease activity index, DAI) were monitored daily and plotted during the induction. (D-F) *Tet2*-deficient mice manifest exacerbated colitis based on colon pathology. Colons from the 4 groups of mice were photographed (D) and quantified based on colon length (E). Damage scores of colons were monitored and plotted based on H&E staining (F). (G) Expression of two classic gut barrier markers *Muc2* and *Occludin* in colon were quantified by qRT-PCR. (H) Damage of gut barrier were quantified by FITC-dextran staining in serum of PB. (I-O) Microbiota (16S rRNA) sequencing analysis of gut in mice feed with DSS or vehicle. The PCA plot of the 4 groups of microbiota samples suggests obvious separation in microbial composition (I). α-diversity of the microbial community based on observed operational taxonomic units were quantified by Simpson or Chao index (J and K). Average relative abundance of prevalent microbiota at the phylum level or at the genus level among the four groups (L and M). Detailed comparison of relative taxon abundance between the *Tet2^+/-^_Veh* vs. *WT_Veh* or between *Tet2^+/-^_DSS* vs. *WT_DSS* are also shown in (N) and (O). PCA, principal component analysis. Data are shown as means ± SEMs in A to H. Number of biological repeats (animals): n = 3∼7. Experiments of DSS treatment on primary mice were repeated three times. *, p < 0.05; **, p < 0.01; ***, p < 0.001; ****, p < 0.0001. See also **Supplemental Figure 2** for representative photography of H&E staining of colons and additional microbiota analysis.

To ask to what extent *Tet2*-defficient primary mice manifest aberrant colon pathogen-host immunity, we performed 16S rRNA sequence and compared the microbial composition between WT and *Tet2*-defficient primary mice (*Tet2^+/-^*_Veh vs. WT_Veh), along with the comparison between WT and *Tet2*-defficient primary mice fed with DSS (*Tet2^+/-^*_DSS vs. WT_DSS). Results of the microbial composition data calculated by principal component analysis (PCA) suggest the four groups distinguish each other very well (**Figure 2I**). Although the Simpson index of operational taxonomic units (OUT, measuring microbial community composition) level fails to distinguish the four groups (a higher Simpson index indicates lower community diversity, **Figure 2J**), the Chaos index of OUT level significantly distinguishes microbial communities of *Tet2^+/-^*_Veh mice from that of WT_Veh mice (a higher Chaos index indicates higher community richness, **Figure 2K**). Upon DSS challenge, the Chaos index of OUT level in *Tet2^+/-^*_DSS and WT_DSS were decreased compared to that in *Tet2^+/-^*_Veh and WT_Veh, suggesting certain bacteria were dominant expanded. Further microbial analysis suggests that: (1) at phylum level, *Tet2*_DSS mice exhibited most increments in *Proteobacteria* and *Actinobacteria* (both are pathogenic and toxic to the animals, **Figure 2L, Supplemental Figure 3A** and **B**); (2) at genus level, different microbiome compositions are revealed for the comparison between *Tet2^+/-^*_Veh and WT_Veh (**Figure 2M**). Detailed comparison between *Tet2^+/-^*_Veh and WT_Veh or between *Tet2^+/-^*_DSS and WT_DSS reveal that aberrant changes in microbial composition including probiotic microbes (circled by blue rectangles) or pathogenic microbes (circled by pink rectangles) (**Figure 2N** and **O**; and **Supplemental Figure 2B** and **C**). Analysis combining hematological parameters and microbial composition further confirm that the pathogenic microbial *Proteobacteria* are involved in alteration of Hemoglobin level and neutrophil percentage (**Supplemental Figure 3C** and **D**). Taken together, results from colon pathology and microbial 16S rRNA sequencing suggest that without DSS feeding, *Tet2*-defficient primary mice have already manifested subtle colitis and aberrant microbial composition, especially increase of pathogenic microbes; upon DSS challenge, *Tet2*-defficient primary mice manifest exacerbated colitis and imbalance of intestinal flora.

### DSS induces aberrant hematopoiesis and skewed myelopoiesis revealed by flow cytometry

To directly clarify the DSS effect on hematopoiesis, cells from blood, spleen and bone marrow were subjected to hematological cell counts, histology, and flow cytometry analysis. Consistently with our previous report that *Tet2*-defficient primary mice manifest skewed myelopoiesis with or without LPS treatment (Cai et al., 2018), we observed similar trends in the *Tet2*-defficient primary mice compared to wild-type controls (**Figure 3A** to **G)**. Flow cytometry results revealed similar skewed myelopoiesis and alterations specific to the pools of HSC and GMP in the *Tet2^+/-^*_Veh or *Tet2^+/-^*_DSS mice compared to the respective controls.

**Figure 3.**
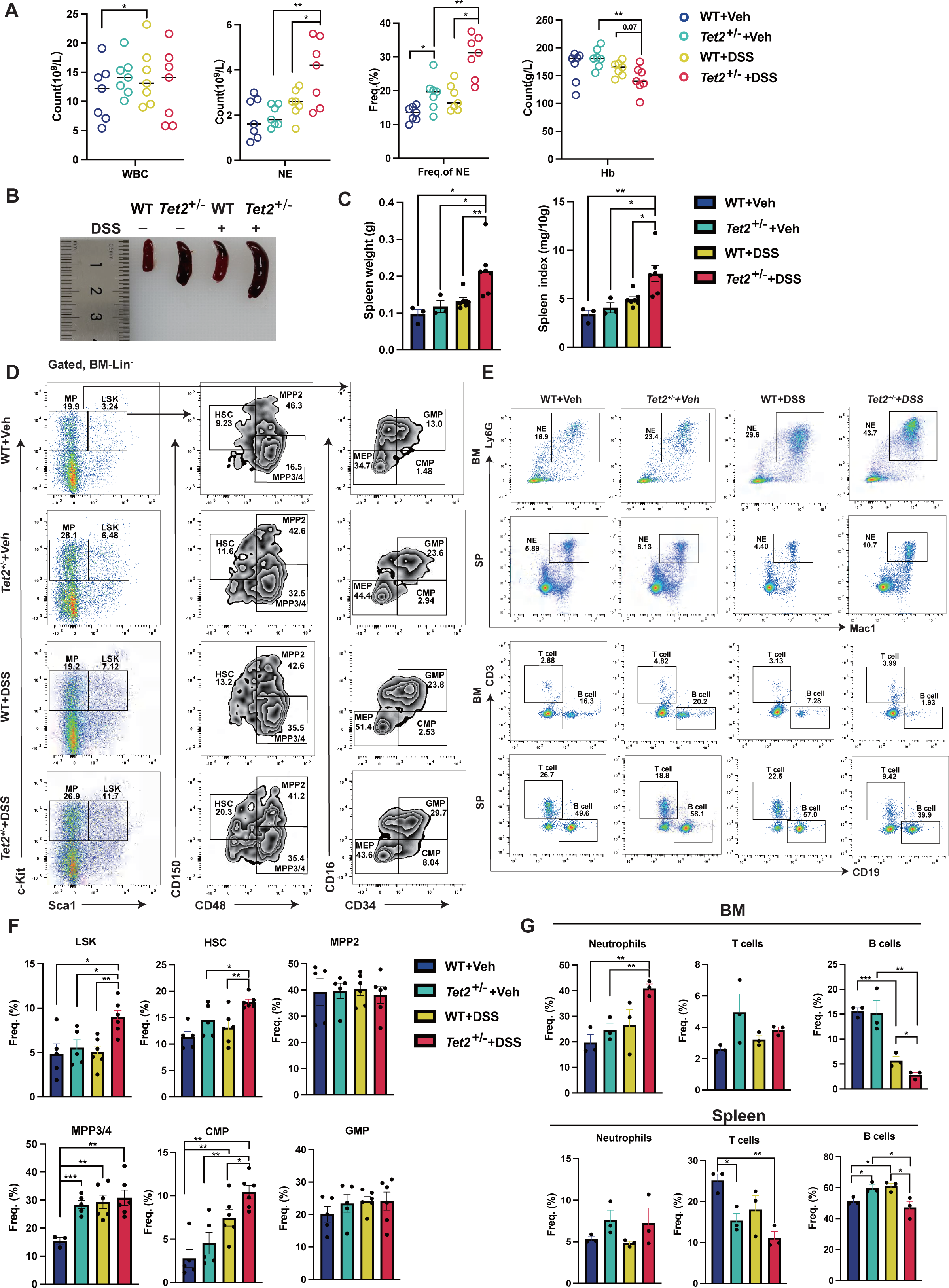
Chronic DSS feeding induces exacerbated myelopoiesis in *Tet2*-deficient mice revealed by flow cytometry. (A) Hematological parameters of PB were monitored at the end-point of the DSS induction. (B and C) Photography and quantification of spleens. (D and F) Gating strategies, representative flow profiles and quantifications of HSPCs in BM. LSK, Lin^-^;Sca1^+^;Kit^+^. HSC, hematopoietic stem cell. MPP, multiple potent progenitor. CMP, common myeloid progenitor. GMP, granulocyte-monocyte progenitor. (E and G) Quantification of mature cells including neutrophils, T cells, and B cells in bone marrow and spleen. BM, bone marrow; SP, spleen; PB, peripheral blood. Data are shown as means ± SEMs. Number of biological repeats (animals): n = 3∼7. Experiments of DSS treatment on primary mice were repeated three times. *, p < 0.05; **, p < 0.01; ***, p < 0.001; ****, p < 0.0001.

### scRNA-seq analysis of colon tissue highlights that cell-to-cell talk between *Ptx3^+^* fibroblasts and *Il-1*β*^+^* monocytes in promoting inflammation

Colon tissue from the four groups of mice was also used for scRNA-seq analysis since this approach can provide a full-map of cell-to-cell talk at the single-cell resolution. After quality controls, around 3000 to 6000 cells from each group were included in the further annotation analysis (**Figure 4A**). A total of 5 main populations including epithelial cells, stromal cells, and immune cells (myeloid cells, B cells, and T cells) are annotated in the UMAP plot (**Figure 4A**). Each group has these five main cell populations. However, *Tet2^+/-^*_Veh, WT_DSS and *Tet2^+/-^*_DSS appear to have a higher percentage of stromal cells compared to the WT_Veh control (**Figure 4B** and **C**). Expression density of the respective markers for the five main cell populations is shown in **Figure 4D**. The overall enriched signaling pathway analysis suggests that inflammatory signaling pathway is upregulated in the myeloid cells of *Tet2^+/-^*_Veh, WT_DSS and *Tet2^+/-^*_DSS compared to the WT_Veh control (green rectangle, **Figure 4E**; and **Supplemental Figure 4A**). Importantly, cell-cycle related pathway is also upregulated in the myeloid cells of *Tet2^+/-^*_Veh compared to WT_Veh control (pink rectangle, **Figure 4E**). Interestingly, we also observed upregulated epithelia-mesenchymal-transition pathway (EMT) in the stromal cells of *Tet2^+/-^*_Veh, WT_DSS and *Tet2^+/-^*_DSS compared to the WT_Veh control (blue rectangle, **Figure 4E**; and **Supplemental Figure 4C**). Overall, the pathway enrichment analysis in each single cell population suggests that myeloid cells and stromal cells grossly have the most alterations in the comparison between *Tet2^+/-^*_Veh vs. WT_Veh or between WT_DSS vs. WT_Veh. We then filtered out myeloid cells and stromal for further cell-to-cell communication.

**Figure 4.**
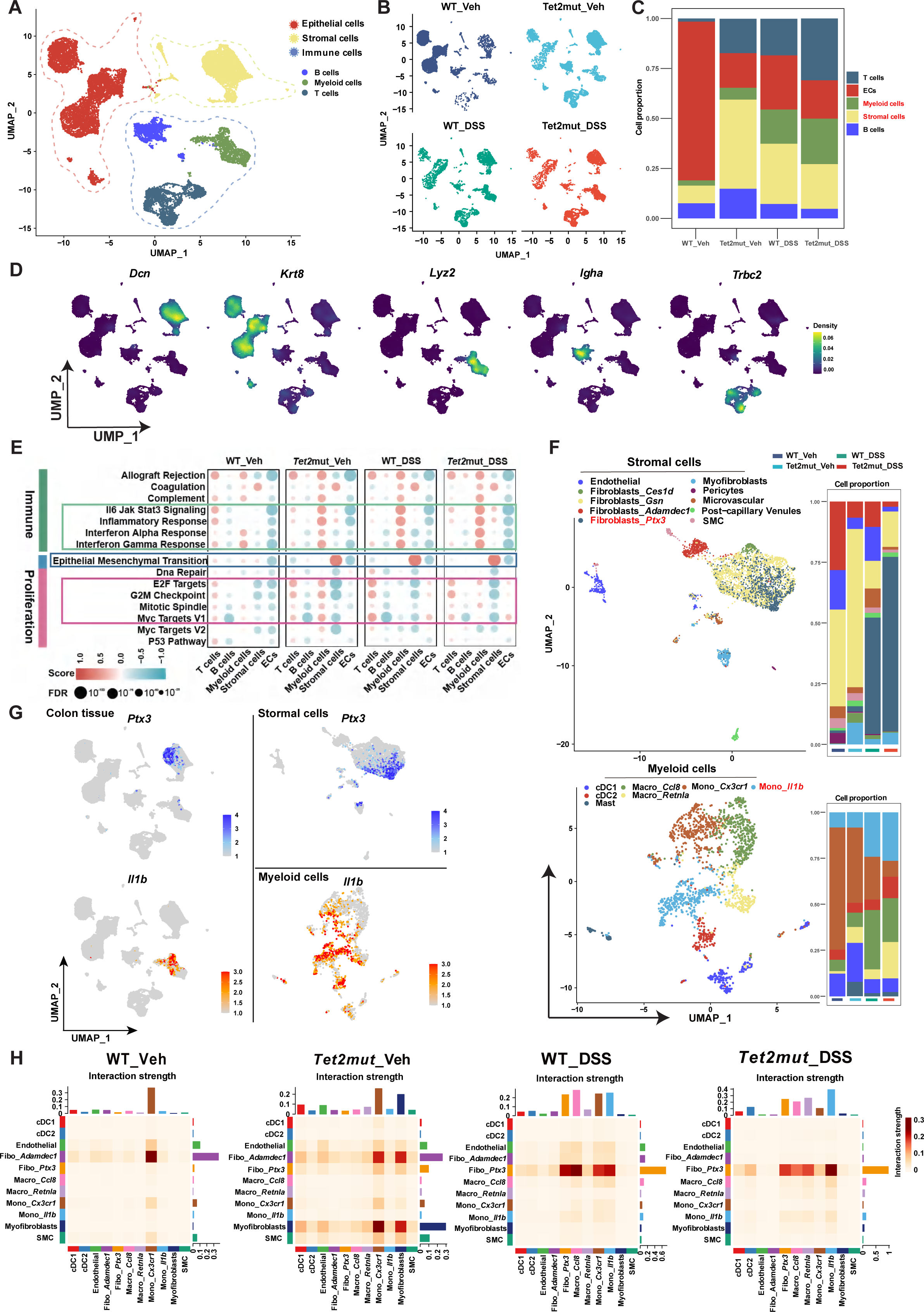
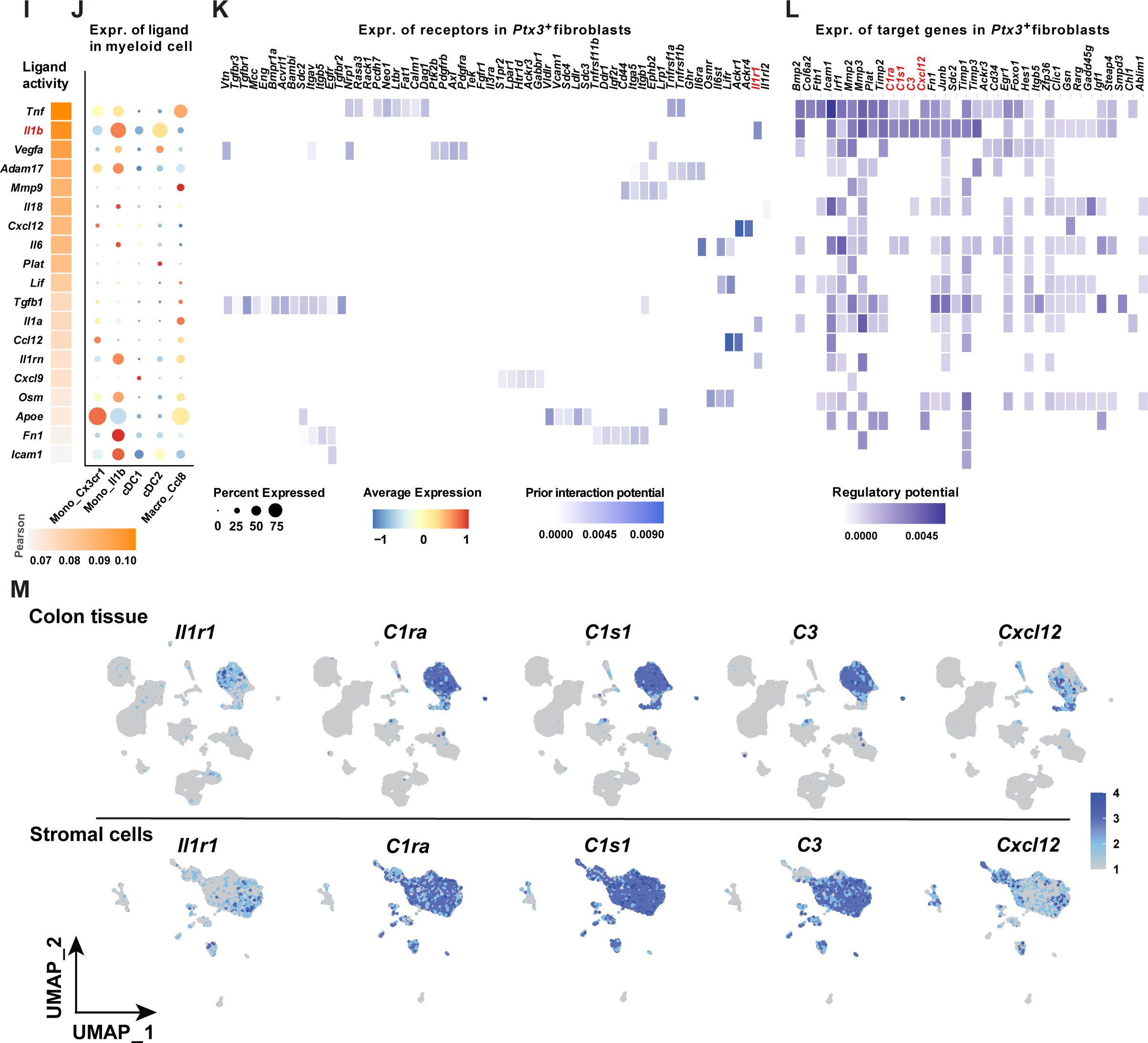
Single-cell RNA-seq analysis of colon tissue from *Tet2*-deficient mice when feed with DSS or vehicle. (A) UMAP plot showing 5 main populations in colon tissues. After lysis, all cells from colon tissues were subjected for scRNA-seq. A total of 18,092 high-quality cells with an average of about 3000 genes per cell were included in the UMAP plot. After quality control during the dataset analysis, the group of *WT_Veh* has 5589 cells; *Tet2mut_Veh* has 5649 cells; *WT_DSS* has 3382 cells; *Tet2mut_DSS* has 4372 cells. The 5 main populations include epithelial cells (red), stromal cells (yellow) and three major types of immune cells: myeloid cells (green), B cells (blue) and T cells (dark blue). (B) Four individual UMAP plots are separately shown according to the sample source. (C) Stacking bar plot showing the portion of the 5 main annotated populations in each colon sample. (D) Expression of representative annotation markers for the 5 main cell populations in the UMAP plot of colons. *Dcn* for stromal cells; *Krt*8 for epithelial cells; *Lyz2* for myeloid cells; *Igha* for B cells; and *Trbc2* for T cells. (E) Heatmap of dysregulated biological pathways in the 5 populations of colon tissues from 4 groups of samples. A full version of the heatmap is shown in the **Supplemental Figure 4**. Of note, Il6-Jak-Stat3 signaling, inflammatory response and interferon alpha response are upregulated in *Tet2mut_Veh, WT_DSS, and Tet2mut_DSS* in myeloid cell, compared to *WT_Veh* (circled by green rectangle). Additionally, proliferation-related pathways appear to be upregulated in myeloid cells of *Tet2mut_Veh* compared to *WT_Veh;* proliferation-related pathways appear to upregulate in T cells of *WT_DSS* and *Tet2mut_DSS* compared to *WT_Veh* (circled by pink rectangle). Interestingly, the epithelial mesenchymal transitional (EMT) pathway appears to be dysregulated in *Tet2mut_Veh, WT_DSS,* and *Tet2mut_DSS* in myeloid cell, compared to *WT_Veh* (circled by blue rectangle). (F) Stromal cells and myeloid cells are subjected for further clustering by UMAP as they have most significantly upregulated biological pathways as indicated in (E). Top panel, UMAP plot of stromal cells and stacking plot showing potion of the sub-population; bottom panel, UMAP plot of myeloid cells and stacking plot showing potion of the sub-population. As indicated, 10 subpopulations and 7 subpopulations are annotated in stromal cells and myeloid cells respectively. (G) Expression of *Ptx3* and *Il-1β* in the colon tissues (left panel) or in stromal cells (up-right panel) or in myeloid cells (bottom-right panel), respectively. (H) Heatmap of cell-to-cell talk strength between subpopulations of stromal cells and myeloid cells. (I-L) Detailed molecular events of the cell-to-cell talk between myeloid cells and stromal cells. Of note, *Ptx3^+^* fibroblasts and *IL-1β ^+^* monocytes appear to be most involved. *IL-1β ^+^* monocytes express numerous ligands including *IL-1β* (I and J). *IL-1β* binds *Il1-1r1* in *Ptx3^+^* fibroblasts and stimulates a couple of downstream genes (K and L). (M) Expression of *IL-1β* receptor *Il1r1* and relevant genes encoding complement components or inflammation regulators (*C1ra*, *C1s1*, *C3* and *Cxcl12*) were plotted on UMAP of colon tissue (top panel) or on the UMAP plot of stromal cells (bottom panel). See also **Supplemental Figure 4** for additional analysis of scRNA-seq colon dataset.

First, stromal cells and myeloid cells from the colon UMAP pool were subject to further clustering to distinguish sub-populations. In total, 8 sub-populations of cells in the stromal cells and 7 sub-populations of cells in the myeloid cells are annotated according to the typical marker gene expression (**Figure 4F**; and **Supplemental Figure 4B**). Expression of *Ptx3* and *Il-1β* is plotted in the UMAP space of colon tissue, stromal cells, or myeloid cells as indicated (**Figure 4G**). To our surprise, expression of *Ptx3* appears to be specific to stromal cells in the colon while expression of *Il-1β* appears to be specific to myeloid cells in the colon in the scRNA-seq dataset. Five sub-populations from the stromal cells and six sub-populations from the myeloid cells had the most alterations in the percentage and therefore were used for calculating cell-to-cell talk strength (**Figure 4H**). Indeed, *Cx3crl1^+^*-myeloid cells appear to have stronger interaction with *Ptx3^+^*-stromal cells in *Tet2^+/-^*_Veh compared to WT_Veh (left two panels, **Figure 4H**). Myofibroblasts also appear to strongly interact with many myeloid cells in *Tet2^+/-^*_Veh compared to WT_Veh (left two panels, **Figure 4H**). In summary, upon DSS feeding, interaction between *Ptx3^+^*-stromal cells and several myeloid cells are dramatically activated (right two panels, **Figure 4H**).

To dissect the detailed molecular events in the *Il-1β ^+^*-myeloid cells and *Ptx3^+^*-stromal cells, five of myeloid sub-populations were used for NicheNet assay (Browaeys et al., 2020). As shown in **Figure 4I** to **L**, IL-1β/IL1-R1 signaling is actively involved in the cell-cell interaction, associated with expression of many downstream genes encoding complement protein proteins or inflammatory ligands (*C1ra*, *C1s1*, *C3* and *Cxcl12*). Surprisingly, *Il1r1* along with *C1ra*, *C1s1*, *C3* and *Cxcl12* all are dominantly expressed in the stromal cells of colon, compared with other four main cell population (**Figure 4M**). Taken together, our detailed cell-cell talk analysis based on the colon scRNA-seq dataset from the 4 groups of mice strongly suggest that PTX3/IL-1β axis recruits inflammatory and complement cascade to module colon immunity in *Tet2*-deficient mice fed with DSS or raised in normal condition.

### Aberrant expression of *PTX3* and *IL-1****β*** are indicated clinical samples with leukemia or colitis

To guarantee that PTX3/IL-1β signaling involves in human colitis or leukemia, we searched publicly available datasets of bulk RNA-seq or scRNA-seq and compared their expression between disease group and healthy controls. Data shown in **Figure 5A** to **D** suggest that increased expression of *PTX3* is detected in the myeloid leukemia group (datasets with AML or MDS patients were analyzed). Interestingly, mutations in *TET2* appear to have higher expression of *PTX3* also (**Figure 5A** and **D**). Importantly, higher expression of *PTX3* has a worse prognosis in the TCGA-LAML cohort while expression of *IL-1*β failed to stratify the cohort (**Figure 5A** and **D**). Additionally, analysis of colitis samples also suggests that higher expression of *PTX3* and *IL-1*β took place in some cohorts of colitis samples compared to the controls (**Figure 5E** to **G**). Taken together, these results warrant that the PTX3/IL-1β bundle indeed involves in human colitis or leukemia.

**Figure 5.**
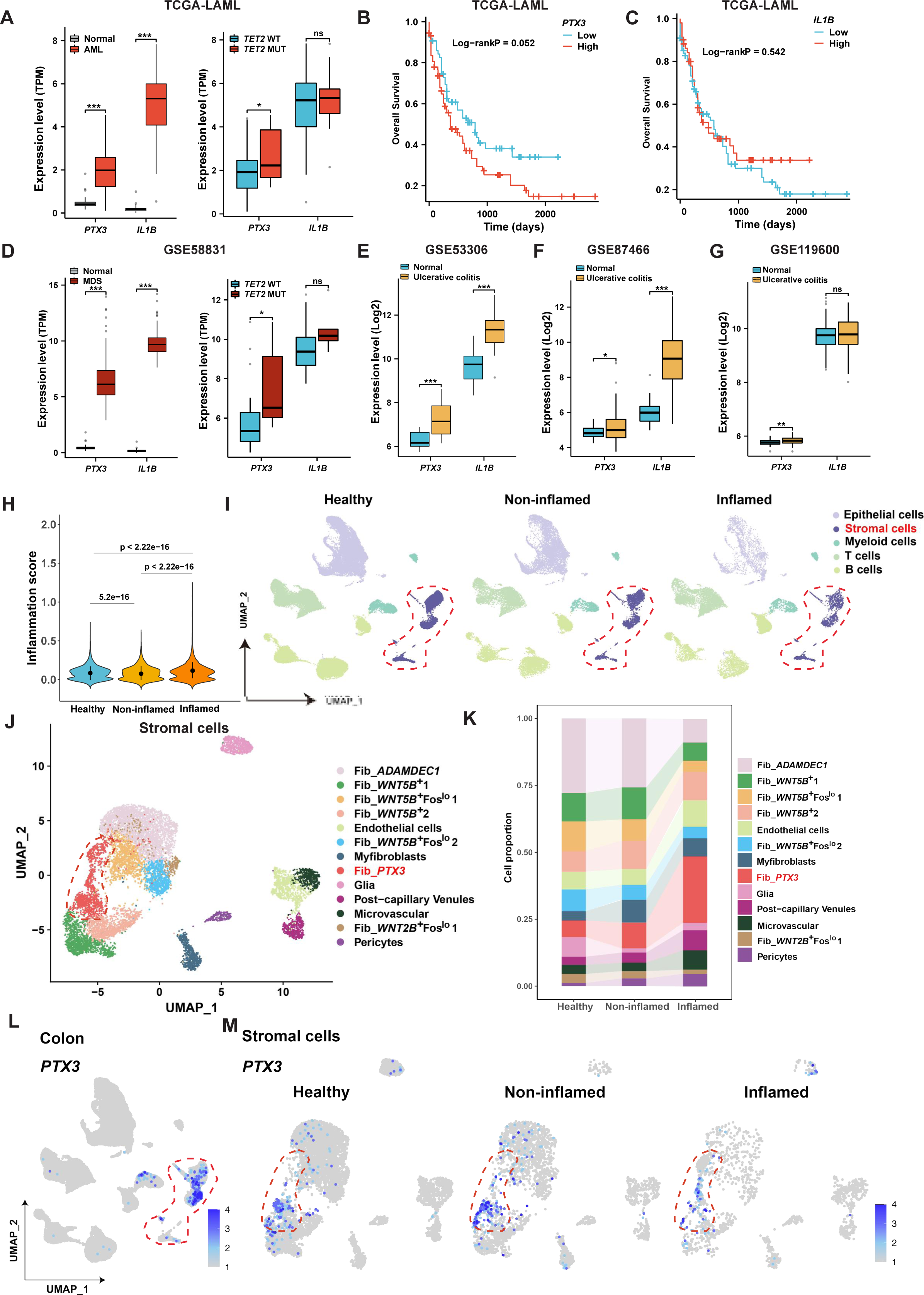
Aberrant PTX3/IL-1β signaling is indicated in human leukemia or colitis through analysis of clinical samples. (A) Expression of *PTX3* and *IL-1*β in unsorted AML patients (left panel) or in AML patients carrying *TET2* mutation (right panel). Bulk RNA-seq datasets of bone marrow were included in the analysis. (B-C) Prognosis value of marker genes *PTX3* and *IL-1*β in the TCGA-LAML cohort. (D) Expression of *PTX3* and *IL-1*β in unsorted MDS patients (left panel, bone marrow samples) or in MDS patients carrying *TET2* mutation (right panel, bone marrow samples). (E-G) Expression of *PTX3* and *IL-1*β in patients with ulcerative colitis and healthy controls, based on three independent cohorts. Bulk RNA-seq datasets of colon tissues were included in E and F; Bulk RNA-seq dataset of blood were included in G. (H-M) Aberrant expression of *PTX3* and *IL-1*β was also detected using scRNA-seq dataset of human colitis samples. The dataset is downloaded from the website Broad Data Use Oversight System (https://duos.broadinstitute.org) and re-analyzed. The cohorts include two groups of human colon samples: colons from healthy controls (n=10) and colons from ulcerative colitis patients (n=7). Patient samples from both non-inflamed regions and inflamed regions were n=7 respectively. Inflammation scores suggest the three groups has significant difference overall (H). UMAP plots of the human colon scRNA-seq dataset also show five main populations as similar as Figure 4A: epithelial cells, stromal cells and three immune cells (myeloid cells, B cells and T cells) (I). Stromal cells were highlighted by the dotted lines (I). Stromal cells were further annotated by UMAP plots and *PTX3^+^*fibroblasts were screened out by dotted lines (J). Of note, stacking plots indicate percentage of *PTX3^+^* fibroblasts was increased according to the inflammation grade of the colitis (K). Expression of *PTX3* is shown in the UMAP plot of colon tissue (L) and three groups of stromal cells (M) as indicated. See also **Supplemental Figure 4** for additional analysis of human colitis scRNA-seq dataset.

As we surprisingly observed that *Ptx3* and *Il-1r1* are dominantly (if not specifically) expressed stromal cells of colon tissue in mice (**Figure 4G** and **M**; and **Supplemental Figure 6A** and **B**), we want to characterize if this pattern is conserved in human. We downloaded the publicly available scRNA-seq datasets of colitis and re-analyzed the cells (Smillie et al., 2019). Results from **Figure 5H** to **M**, along with **Supplemental Figure 5,** suggest that the percentage of *PTX3^+^*-fibroblasts indeed increased according to the grade of colitis. Although partial myeloid cells express *PTX3* themselves, *PTX3* is also dominantly expressed in stromal cells. Taken together, results from human clinical samples suggest that PTX3/IL-1β signaling may have important functions in colitis and leukemia.

### Pharmacological blockage of IL-1**β** signaling by Anakinra inhibits PTX3 expression and inflammation in colon and mitigates TedCH in blood

PTX3 is a soluble pattern recognition molecule (PRM) and has been reported to be directly involved in IL-1β signaling (Bonavita et al., 2015). We postulate that the PTX3/IL-1β signaling pathway is present at the protein level in the colon of *Tet2^+/-^* mice with chronic inflammation. To validate this hypothesis, we performed western blot analysis to examine the expression of Il-1β and Ptx3 (**Supplemental Figure 6C** and **D**). Additionally, immunohistochemical (IHC) staining revealed high levels of Ptx3 expression in inflammatory cells infiltrating the colonic tissues of *Tet2^+/-^* mice (**Supplemental Figure 6E** and **F**).

As inhibitors specific to PTX3/Ptx3 activity are not available, we turn to use IL1R1 specific inhibitor Anakinra to assess the function of Ptx3 in the TedCH, colitis and aberrant hematopoiesis. cBMT assays for TedCH were conducted along with DSS and/or Anakinra treatment (**Figure 6A**). Consistent with its anti-inflammation function, application of Anakinra mitigate TedCH expansion in the mice feed with Vehicle or DSS (**Figure 6B**; and **Supplemental Figure 6G**). Aberrant hematopoiesis also appears to be partially rescued in the TedCH_Anakinra mice compared to TedCH_PBS mice (**Figure 6B**; and **Supplemental Figure 6H** and **K**). Importantly, application of Anakinra reduced the expression of both Ptx3 and Il-1β (**Figure 6D** and **E**). The reduced expression of Ptx3 upon Anakinra is also validated by immunohistochemistry (IHC); pathologic index is also largely ameliorated upon Anakinra treatment (**Figure 6F** to **I**). Taken together, results from Anakinra treatment suggest that inhibition of Il-1β signaling modulated Ptx3 expression and inflammation severity, along with TedCH trajectory and aberrant hematopoiesis.

**Figure 6.**
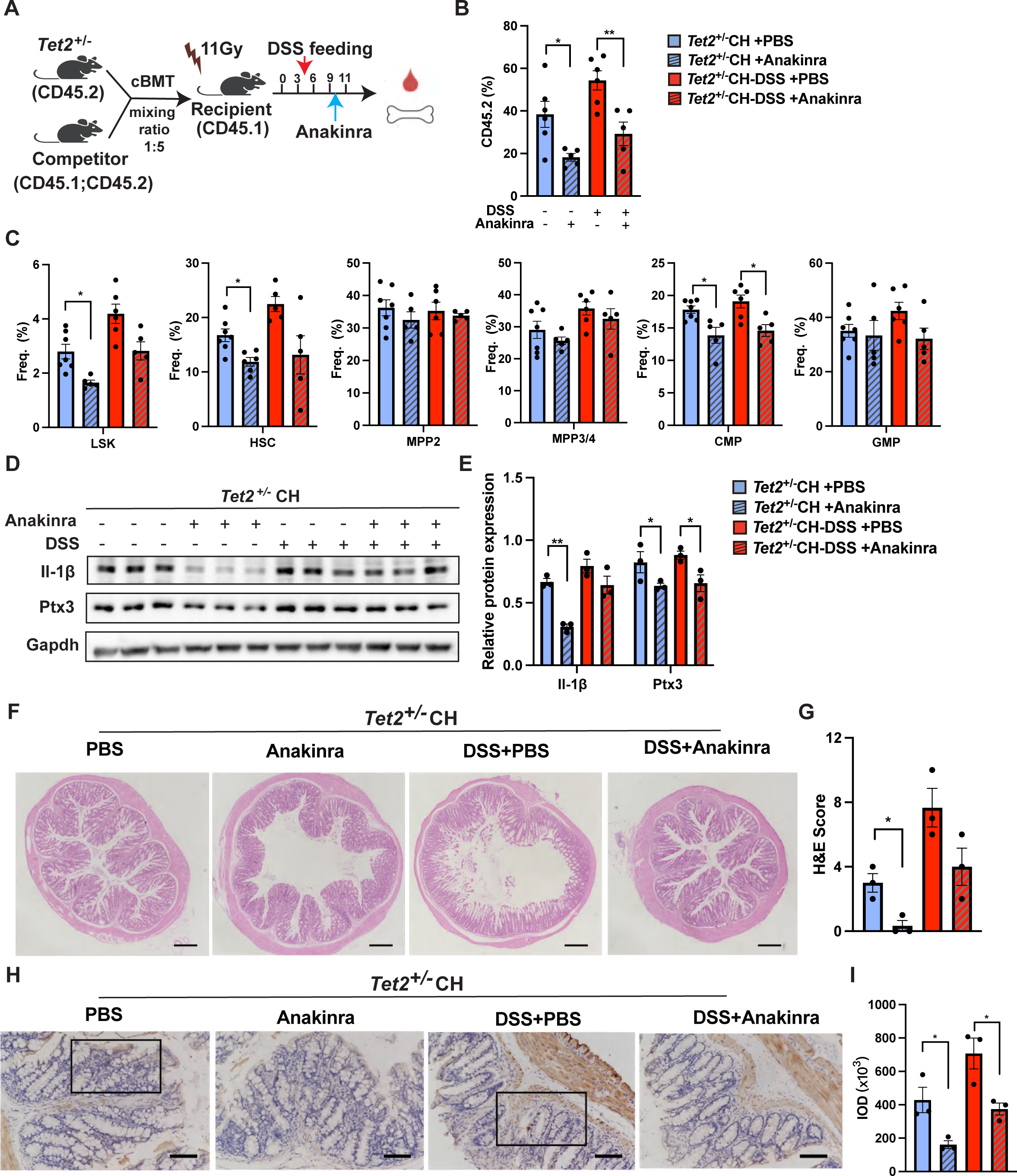
Pharmacological blockage of IL-1β signaling by Anakinra inhibits Ptx3 expression and inflammation in colon and mitigates TedCH in blood. (A) Schematic representation of Anakinra treatment on the DSS-induced expedited TedCH. (B) Quantification of PB chimerism in TedCH mice treated with DSS and/or Anakinra. (C) Quantification of HSPCs including LSK, HSC, MPP2, MPP3/4, CMP and GMP in 4 groups of TedCH mice by flow cytometry. (D-E) The protein levels of IL-1β and Ptx3 were measured in colon tissues by western blot and quantified. (F-G) Representative H&E staining of colon tissue from each group and disease scores (damages in colon tissue) were quantified accordingly. Scale bar is 500 μm. (H-I) Representative immunohistochemical (IHC) staining of Ptx3 in colon (H) and semi-quantitative analysis of immunohistochemistry staining of Ptx3 based on integrated optical density (IOD) (I). scale bar is 100 μm. Data are shown as means ± SEMs in A-H. Number of biological repeats (animal mouse): n = 5∼7. Experiments of DSS treatment combined with Anakina on TedCH mice were repeated twice. *, p < 0.05; **, p < 0.01; ***, p < 0.001; ****, p < 0.0001. See also **Supplemental Figure 4** for additional analysis of Ptx3 expression in TedCH colons.

**Figure 7.**
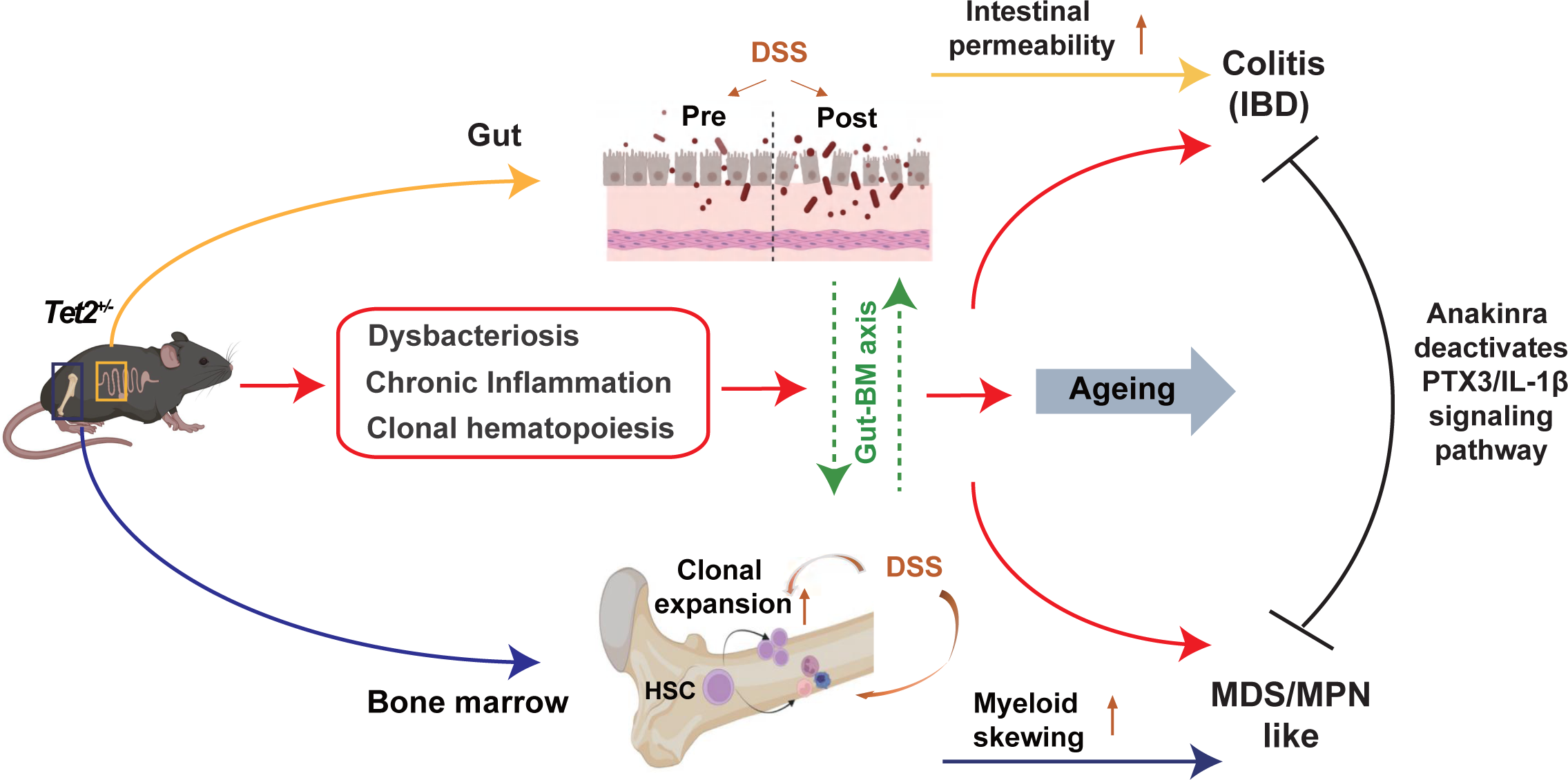
Schematic illustration for the working model about clonal hematopoiesis cooperating with inflammation and dysbacteriosis in gut-bone marrow axis to promote colitis and leukemia. We propose that the gut-bone marrow axis plays an important role in the co-symptoms of colitis and leukemia in the *Tet2*-mutant chimeric and primary mice. Dysbacteriosis, chronic inflammation and clonal hematopoiesis are all assembled to promote the progression the co-symptoms over age. The PTX3/IL-1b signaling pathway plays an important role in promoting colitis and leukemia. See the main text for further discussion.

## Discussion

Since clonal hematopoiesis was for the first time recognized as a new risk medical entity to broadly predict diseases and even all-cause mortality around a decade ago, numerous genetic drivers or passengers of clonal hematopoiesis have been characterized, along with a recent report indicating their trajectories were able to be mathematically computed in human samples (Abelson et al., 2018; Fabre et al., 2022; Genovese et al., 2014; Weeks et al., 2022; Weinstock et al., 2023).To understand the mechanisms underlying CH, it was proposed that different genetic drivers of CH may favor corresponding environmental factors to strengthen or amplify the onset and consequence of CH (Caiado et al., 2021). For example, mutations in *TP53* and *PPM1D,* two top-10 prevalent drivers of CH, were selected when irradiation was applied in the therapy or after exposure to cisplatin and doxorubicin treatment (Wong et al., 2015) (Hsu et al., 2018). Our previous studies have shown that mutations in *TET2*, the first identified and one of the top-3 driver of CH, favors inflammatory condition on the contrary; blocking inflammation by genetic loss of *Morrbid* or administration of anti-inflammation drugs E3330 or SHP099 mitigate TedCH in experimental models (Cai et al., 2020a; Cai et al., 2018; Cai et al., 2021; Cai et al., 2020b). However, whether and how other environmental factors (positive, negative, or neutral to TedCH), intra-cell or intra-organ or host-pathogen communications drive TedCH remain incompletely understood.

To reach these goals, we first assessed the role of five different environmental factors in TedCH (4 factors are included in this study; and another one based on *Tet2^+/-^* HSPCs mixed with genetically stable inflammatory HSPCs will be reported elsewhere) and prioritized the impact of DSS, an inducer of infection with toxic pathogens and colitis in gut. We further characterized the impact of DSS on the *Tet2*-deficient chimeric or primary mice by multi-omic tools including single cell RNA sequencing or gut microbe sequencing. Although administration of the other three challenges, irradiation, 5-FU (depletion of immune cells and hematopoietic cells), or STZ (toxic damager of pancreases and inducer of high blood glucose) failed to accelerate TedCH, chronic colitis induced by three cycles of feeding with DSS or mixing with IL-1β-secreting genetically stable inflammatory HSPCs expedite the trajectory of TedCH. These results suggest that: 1) only certain specific environmental factors play a positive role to cooperate with TedCH in experimental models; 2) these “positive” factors may have a direct link to inflammation or innate immunity; 3) the gut-bone marrow axis, which could be stimulated by host-pathogen axis or just by dysregulated autoimmunity, play a role in TedCH and leukemogenesis.

These observations from us are consistent with a previous study by *Meisel* et al. who directly studied the pathogen-host interaction in *Tet2*-mutation mediated leukemic diseases. In the study, *Meisel* et al. studied why *Tet2*-mutant primary mice did not develop MPN or CMML phenotypes with 100% penetration and proposed that microbial signals cooperate to induce pre-leukemic MPN phenotype in the mice (host-pathogen axis in TedCH-related diseases). Using *Tet2*-mutant mice raised in germ-free conditions, the authors further validate their conclusions (Meisel et al., 2018; Zeng et al., 2019). However, most of the experimental mice and our human are exposed to microbial environments during life. Where the initial inflammation takes place (gut, bone marrow or even other organ?) and how it triggers CH and changes its trajectory remain to be elucidated.

Considering the importance of inflammation in *Tet2*-mutation related diseases, in previous studies we integrated multiplayers and proposed a working model for *Tet2*-mutation related diseases: TedCH recruits a feed-forward loop and inflammation acts as an amplifier in the loop for disease progression including chronic leukemia and acute leukemia (Cai et al., 2018; Cai et al., 2021). Given that DSS promotes TedCH in chimeric mice, we then tested if DSS could promote *Tet2-*mediated CMML to transform into full-blown AML. During the short-period of colitis, we indeed notice exacerbated phenotypes in *Tet2*mutant_DSS mice in both gut permeability (worsen colitis) and bone marrow hematopoiesis (malignant transformation to MDS/MPN-like). Unlike that we observed full-blown AML in aged *Tet2^+/-^; Flt3^ITD/ITD^* or *Tet2^+/-^;Ins2^+/-^*, we failed to observe full-blown AML in the DSS-challenged mice. These results suggest that inflammation as an amplifier to induce full-blown AML may depend on adequate time.

Furthermore, single-cell RNA-seq (scRNA-seq) today has become a very powerful tool to dissect the etiology of complicated diseases at the single cell level and molecular level. Both scRNA-seq and bulk RNA-seq were applied in this study to delineate the widespread transcriptional changes in gut. Analysis of scRNA-seq in gut of *Tet2*-mutant_Veh and *Tet2*-mutant_DSS mice helps us to reveal a previously unappreciated alteration in *Ptx3^+^*-fibroblast of gut and the inferred PTX3/IL-1β signaling [note that mice were raised in specific pathogen-free (SPF) condition with or without feeding with DSS]. These results make us conclude that *Tet2*-mutant mice appear to develop colitis and leukemia simultaneously, very subtle if not strong, reminiscent of human symptoms. Consistently, the involvement of PTX3/IL-1β inflammatory signaling is also confirmed when we re-analyzed human colitis scRNA datasets.

Interestingly, recent clinical studies have also shown that patients with inflammatory bowel disease (IBD) are associated with clonal hematopoiesis and mutations in *DNMT3A* and *PPM1D* are prevalent although mutations in *TET2* are not dominant in the cohort on the contrary (Feng et al., 2023; Zhang et al., 2019). The colitis patients harbor mutant hematopoietic clones and often present with clinical symptoms of MDS/AML (Cumbo et al., 2022). Moreover, intestinal barrier dysfunction is also reported at diagnosis in patients with MDS and AML (Khan et al., 2021). Based on the present study, along with these clinical observations, we may conclude that *Tet2*-mutant mice manifest co-symptoms of colitis and leukemia and that a molecular link, PTX3/IL-1β signaling, modulate both diseases in the mice and in human. Further clinical samples with mutations in various clonal hematopoiesis drivers will help answer these questions.

Changes in the gut microbiome are considered a key regulator of the conversion of normal hematopoiesis to diseased hematopoiesis, such as anemia and neutropenia caused by IBD or long-term antibiotic use (Deshmukh et al., 2014). A study has shown that an imbalance in the microbiota induced by antibiotic exposure or sterile conditions can lead to dysregulation of hematopoiesis (Yan et al., 2018). Our data from 16S rRNA gene sequencing indicated that *Tet2^+/-^* mice had disturbed gut microbiota with significantly reduced probiotic abundance prior to the addition of inflammatory factors. For example, chronically infected *Tet2^+/-^* mice exhibited increased *Proteobacteria* and *Actinobacteria*, two toxic gut pathogens to mammals. We also observed that there was a positive correlation between these microbiotas and PB hematological parameters (**Supplemental Figure 3C** and **D**). Previous clinical studies have shown that patients undergoing hematopoietic stem cell transplantation (HSCT) have reduced gut microbiota diversity and increased *Proteobacteria*, which correlates with the risk of patients developing bloodstream infections during HSCT (Taur et al., 2012). These results suggest a potential role of microbial signaling in exacerbating the disease burden of IBD in *Tet2^+/-^* mice as well as hematologic malignancy transformation. Functional analysis in the appropriate facility with adequate biological protection, for example, pathogen infection or fecal microbial transplantation, will offer direct evidence with regards to the interaction between specific-pathogens and TedCH or *Tet2*-mutant mice (Zeng et al., 2023).

Consistent with previous studies (Burns et al., 2022; Caiado et al., 2023), our data suggest that Anakinra eliminates the *Tet2^+/-^* clones in neutrophil and HSPC compartments, inhibits gut inflammation, restores the adaptive hematopoietic injury induced by chronic inflammation, and attenuates colitis. Dissecting the role of *PTX3* and *IL1R1* in a cell-type specific deletion will assist in further understanding the cellular communications, i.e., immune cells with epithelia cells and offer a new knowledge for understanding gut-bone marrow axis and co-symptoms of colitis and leukemia. Based on our pharmacological data using the IL-1R1 inhibitor Anakinra, it is feasible to speculate that: 1) cocktails of antibiotics administration would also be able to slowdown TedCH, which should be tested in future studies; 2) blocking the IL-1β/PTX3 axis may represent a promising therapeutic target for preventing and treating infection-related clonal hematopoiesis and associated diseases. Since specific PTX3 inhibitors, except PTX3 antibodies, are still not available in research-grade or clinical-grade, developing PTX3 inhibitors or combining them with methylation drugs may represent potential therapeutic approaches for targeting *TET2* mutant hematopoiesis in the future.

In summary, the present study demonstrates that gut-bone marrow axis plays an important role in the co-symptoms of colitis and leukemia in the *Tet2*-mutant chimeric and primary mice. Dysbacteriosis, chronic inflammation and clonal hematopoiesis are all assembled to promote the progression of the co-symptoms over time. Our single-cell transcriptomic analysis highlights and appreciates the role of a population of gut *Ptx3^+^*-fibroblasts in driving the entire diseases. Mitigation of inflammation by IL-1R1 inhibitor Anakinra can downregulate Ptx3, repair gut barrier and slowdown TedCH. The study therefore demonstrates that PTX3/IL-1β signaling-associated clonal hematopoiesis plays important roles in the gut-bone marrow axis and related chronic diseases, including colitis and leukemia.

## Acknowledgments

We thank members of Cai laboratory and colleagues of Tianjin Medical University for their technical and administration support, as well as for their helpful suggestions improve the manuscript. We would also like to thank Drs. Mingjiang Xu, Tao Cheng, Hui Cheng, Zhiqiang Liu, and Mi Deng for sharing us reagents.

## Author Contributions

ZC and HH conceived and designed the study, wrote the computational scripts for sequencing analysis, visualized the results, drafted, and wrote the manuscript. HH and YW performed most of the experiments and analysis. HY, QH, WJ, ZW, GD and JD assisted the experimental verification or validations from other database/cohorts and edited the manuscript. HW and ZZ contributed critical reagents and analyzed the data. All authors contributed to the editing and revision of the manuscript.

## DECLARATION OF INTERESTS

ZC is a scientific advisor to Beijing SeekGene BioSciences Co. Ltd. GD is an employee of Beijing SeekGene BioSciences Co. Ltd. Other authors declare no potential conflict of interest.

## Funding

This work was supported in part by grants from the Tianjin Medical University Talent Program and from National Science Foundation of China to ZC (No.82170173, No. 82371789).

## STAR METHODS

### KEY RESOURCES TABLE

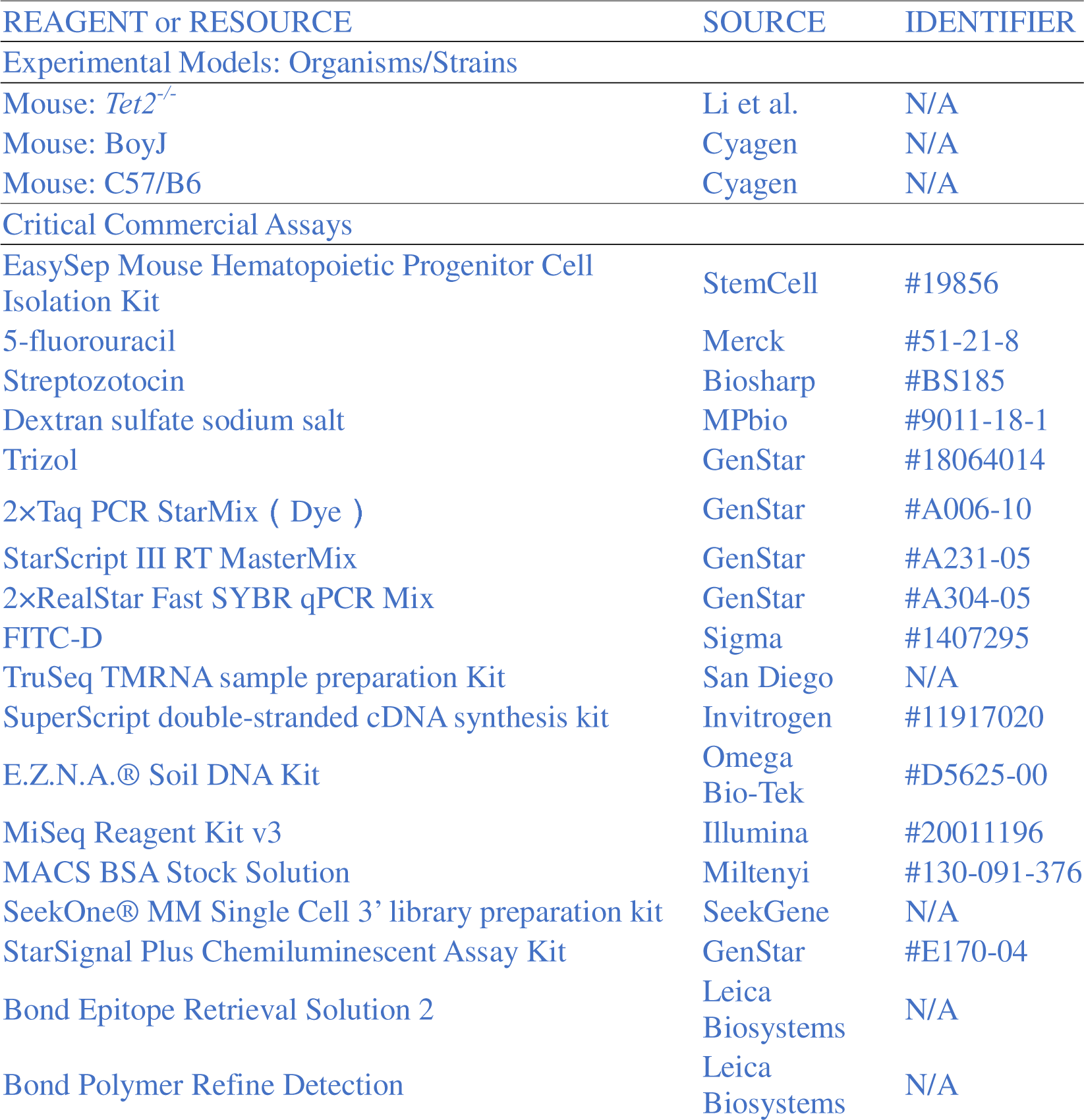
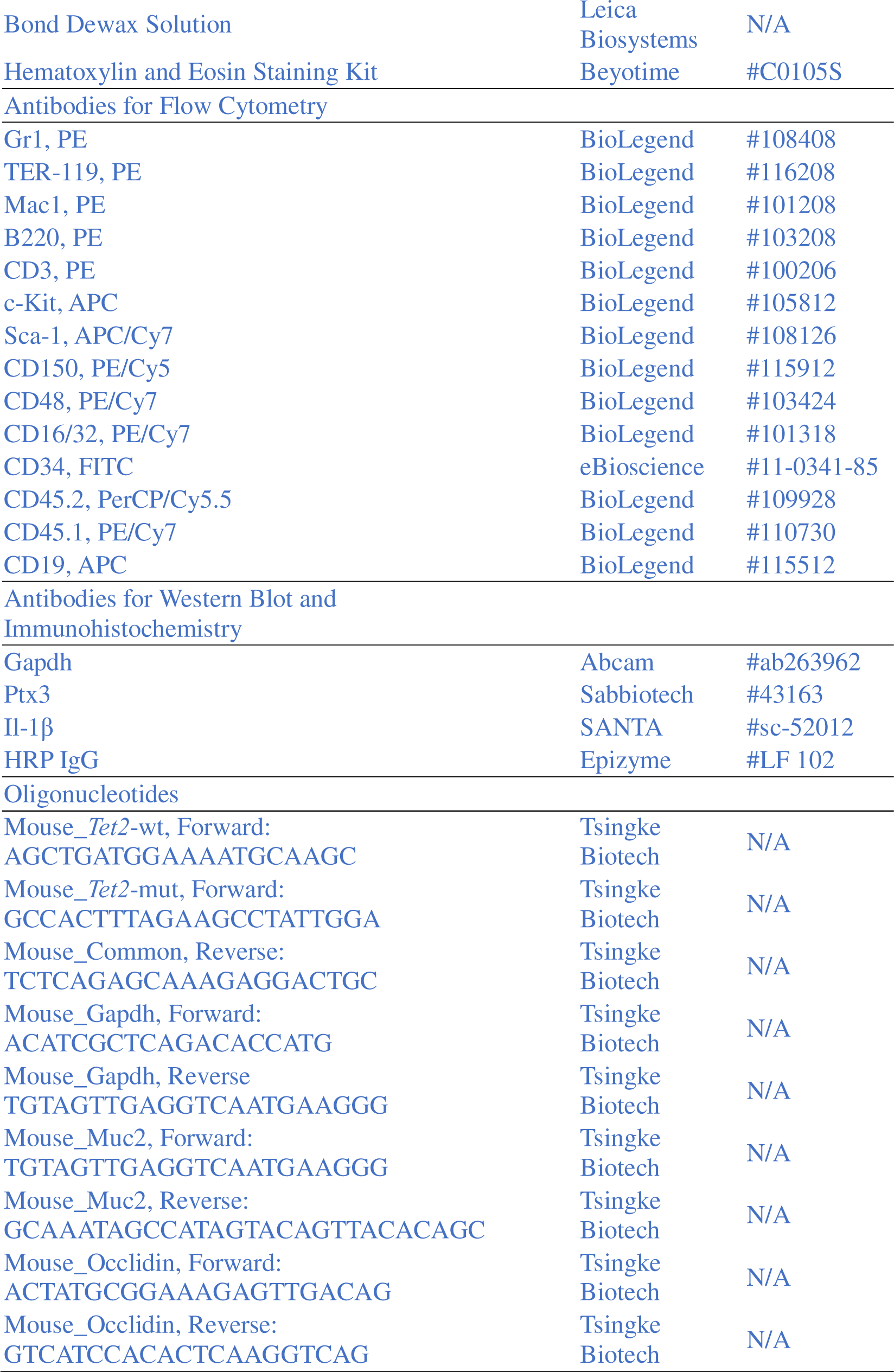

## RESOURCE AVAILABILITY

### Lead contact example text

Further information and requests for resources and reagents should be directed to and will be fulfilled by the lead contact, Zhigang Cai.

### Materials availability

This study did not generate new unique reagents.

### Data and code availability

Any additional information required to reanalyze the data reported in this paper is available from the lead contact upon request.

The raw sequencing data from this study have been deposited in the Genome Sequence Archive in BIG Data Center (https://bigd.big.ac.cn/), Beijing Institute of Genomics (BIG), Chinese Academy of Sciences, under the accession number: PRJCA016651 (the datasets will be available to the public once the manuscript in press).

This paper does not report any original code.

## EXPERIMENTAL MODEL AND STUDY PARTICIPANT DETAILS

### Mice

We used wild-type C57BL/6 (CD45.2), C57BL/6.SJL (CD45.1, BoyJ) and *Tet2^+/-^* mice at the age of 8 weeks (Li et al., 2011). Animal experimentation was performed in accordance with protocols approved by the Animal Care and Use Committee of Tianjin Medical University.

## METHOD DETAILS

### Competitive bone marrow transplantation and In vivo assays

Recipient animals (F1, CD45.2/ CD45.1) were lethally irradiated (7Gy plus 4Gy) one day prior to transplantation (intravenous tail injection) of donor cells. For generating chimeric mice mimicking hematopoietic clonal expansion, *Tet2^+/-^* donor cells and BoyJ donor cells were mixed at a ratio of 1: 5 (100K: 500K). 4 weeks after BM transplantation animals were injected with 5-FU (5-fluorouracil F6627, Sigma-Aldrich) at a dose of 150 mg/kg in 100ul of PBS. Mice were given a single intraperitoneal injection with a dose of 180 mg/kg STZ (streptozocin, BS1485, Biosharp) after fasting for more than 12 hours to establish hyperglycemia model. Mice were treated with ddW with or without 2.5% DSS (dextran sulfate sodium, MP Biomedicals) for 1 week, ddW for another week for three cycles. For pharmacological assays, Anakinra (GC39339, GLPBIO) was injected intra-peritoneally at a dose of 37ug/mouse in 100ul of PBS every other day for 1 month.

### Flow cytometry

The BM cells were flushed out with FACS buffer. Single-cell suspensions were treated with red blood cell lysis buffer, stained, and analyzed using FACS Canto II (BD Biosciences). For mature cell analysis, staining with antibodies against B220 (#103208, BioLegend), CD3 (#100206, BioLegend), Gr-1 (#108408, BioLegend), CD11b (#101208, BioLegend). For early hematopoietic cell analysis, cells were incubated with biotinylated antibodies against the lineage (Lin) markers: CD11b, Ter119 (#116208, BioLegend), Gr-1, CD3, B220, and the fluorescence-conjugated antibodies: c-Kit (#105812, BioLegend), Sca-1 (#108126, BioLegend), CD34 (#11-0341-85, BioLegend), CD150 (#115912, BioLegend), CD48 (#103424, BioLegend), and CD16/32 (#101318, BioLegend).

### Single cell RNA-sequencing

Immature hematopoietic cells (Linage-negative, Lin^-^) was collected by EasySep^TM^ cell separation kit (Stem Cell Co.). After harvested, cell count and viability of the bone marrow were estimated using fluorescence Cell Analyzer (Countstar^®^ Rigel S2) with AO/PI reagent after removal erythrocytes (R1010, Solarbio) and then dead cells removal was decided to be performed or not (130-090-101, Miltenyi). Finally fresh cells were washed twice in the RPMI1640 and then resuspended at 1×10^6^ cells per ml in 1×PBS and 0.04% bovine serum albumin.

Colon tissues were washed in ice-cold RPMI1640 and dissociated using Collagenase □ (V900891-100MG, Sigma) and Collagenase □ (C5138-500MG, Sigma). DNase □ (9003-98-9, Sigma) treatment was optional according to the viscosity of the homogenate. Cell count and viability was estimated using fluorescence Cell Analyzer (Countstar^®^ Rigel S2) with AO/PI reagent after removal erythrocytes (R1010, Solarbio) and then dead cells removal was decided to be performed or not (130-090-101, Miltenyi). Finally fresh cells were washed twice in the RPMI1640 and then resuspended at 1×10^6^ cells per ml in 1×PBS and 0.04% bovine serum albumin.

Single-cell RNA-Seq libraries were prepared using SeekOne^®^ MM Single Cell 3’ library preparation kit (No.K00104, SeekGene). Briefly, the appropriate number of cells were loaded into the flow channel of SeekOne^®^ MM chip which had 170,000 microwells and allowed to settle in microwells by gravity. After removing the unsettled cells, sufficient <u>C</u>ell <u>B</u>arcoded Magnetic <u>B</u>eads (CBBs) were pipetted into flow channel and also allowed to settle in microwells with the help of a magnetic field. Next excess CBBs were rinsed out and cells in MM chip were lysed to release RNA which was captured by the CBB in the same microwell. Then all CBBs were collected and reverse transcription were performed at 37□ for 30 minutes to label cDNA with cell barcode on the beads. Further Exonuclease I treatment were performed to remove unused primer on CBBs. Subsequently, barcoded cDNA on the CBBs was hybridized with random primer which had reads 2 SeqPrimer sequence on the 5’ end and could extend to form the second strand DNA with cell barcode on the 3’ end. The resulting second strand DNA were denatured off the CBBs, purified and amplified in PCR reaction. The amplified cDNA product was then cleaned to remove unwanted fragments and added to full length sequencing adapter and sample index by indexed PCR. The indexed sequencing libraries were cleanup with SPRI beads, quantified by quantitative PCR (KK4824, KAPA Biosystems) and then sequenced on illumina NovaSeq 6000 with PE150 read length.

### Fecal microbiota evaluation

Mice were individually placed in clean cages for feces collection. Fresh fecal samples were collected into sterile cryopreservation tubes, quickly frozen in liquid nitrogen and stored at −80°C. Purified amplicons were pooled in equimolar and paired-end sequenced on an Illumina MiSeq PE300 platform/NovaSeq PE250 platform (Illumina, San Diego,USA) according to the standard protocols by Majorbio Bio-Pharm Technology Co. Ltd. (Shanghai, China).

### H&E Staining and Immunohistochemistry Staining

The tissue samples were fixed in 4% paraformaldehyde. After the dehydration in ethanol, the tissue was embedded in para□n then sectioned with a thickness of 4 μm. The histopathological feature was tested via H&E staining. To perform immunohistochemistry staining, the tissue sections were depara□nized, rehydrated, and rinsed, followed by antigen retrieval and blocking (goat serum). Next, the tissue sections were incubated with primary antibody overnight and followed by the biotinylated secondary antibodies. The DAB Horseradish Peroxidase Color Development Kit (Dako, Agilent Technologies, USA) was applied for color reaction in immunohistochemistry staining. Finally, the tissue sections were observed under the optical microscope or fluorescence microscope. The Image J software was applied for scoring the immunohistochemistry picture.

Immunohistochemical (IHC) staining was using a Leica Bond RX stainer (Leica, Buffalo Grove, IL). Slides were retrieved for 20 min using Epitope Retrieval 1 (Citrate; Leica) and incubated in Protein Block (Dako, Agilent, Santa Clara, CA) for 5 min. Primary antibodies were diluted in Background Reducing Diluent (Dako) as follows: Ptx3 (41372, Sabbiotech, USA) at 1:900, which was diluted in Bond Diluent (Leica) at 1:200. Primary antibodie was diluted in Background Reducing Diluent (Dako) and incubated for 15 min. Immunostaining visualization was achieved by incubating slides 10 min in DAB and DAB buffer from the Bond Polymer Refine Detection System. Slides were counterstained for 5 min using Schmidt hematoxylin, followed by several rinses in 1x Bond wash buffer and distilled water.

### Western blot analysis

Proteins were harvested from cells and colon tissues with RIPA Lysis Buffer (Beyotime, Jiangsu, China) supplemented with phenylmethyl sulfonyl fluoride (PMSF) protease inhibitor and phosphatase inhibitor. Total protein concentration was determined by BCA Protein Assay Kit (Beyotime, Jiangsu, China), denatured protein samples of appropriate quality of proteins were subjected to sodium dodecyl sulfate polyacrylamide gel electrophoresis (SDS-PAGE) and then transferred to PVDF membranes. Then membranes were later blocked with 5% skimmed milk, and incubated were immunodetected with specific antibodies against Il-1β (515598; SANTA, USA), Ptx3 (41372, Sabbiotech, USA), and GAPDH (ab181620, Abcam, USA) overnight at 4 °C. Protein bands were visualized by the MINICHEMI Imaging System (Surwit, Hangzhou, China) using the commercial Pierce ™ Fast Western Blot Kit and the ECL Substrate (GenStar, Beijing, China).

### List of computation & visualizing software for mouse scRNA-seq

Most of the software used for computing and visualizing the scRNA-seq of both mouse and human have been described in our previous study (He et al., 2022). Coding scripts is available upon request.

## QUANTIFICATION AND STATISTICAL ANALYSIS

### Statistical analysis

Statistical analysis was conducted by using GraphPad 9. Comparisons between 2 groups were determined by using a two-tail student’s t-test. Comparison of multiple groups were determined by using an ANOVA analysis of variance with the Dunnett multiple comparisons test. Most of experiments in this study were repeated 2 or 3 times independently and representative data was shown. Results with p < 0.05 were considered statistically significant.

---

**Supplementary Figure 1.**
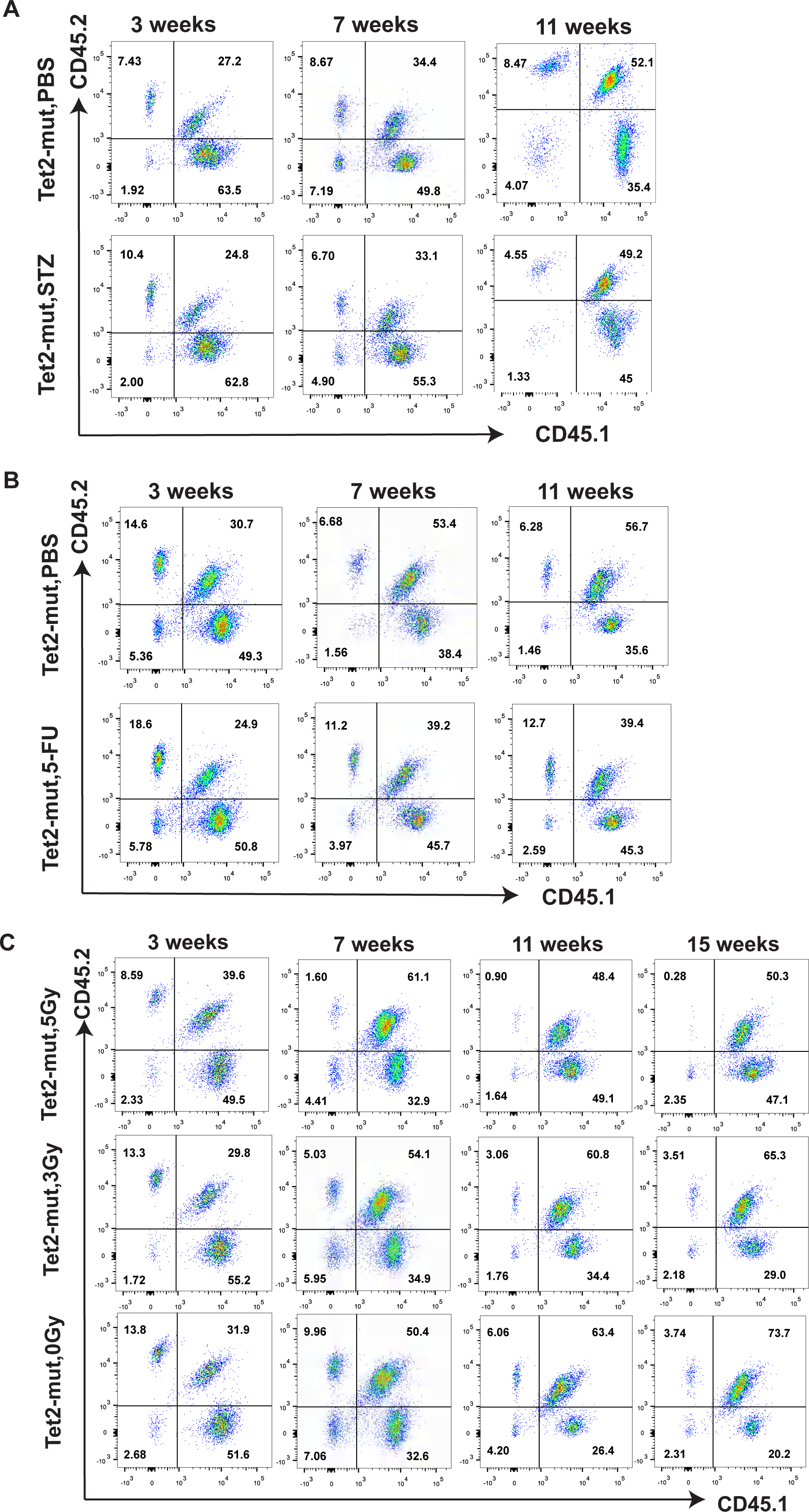

**Supplementary Figure 2.**
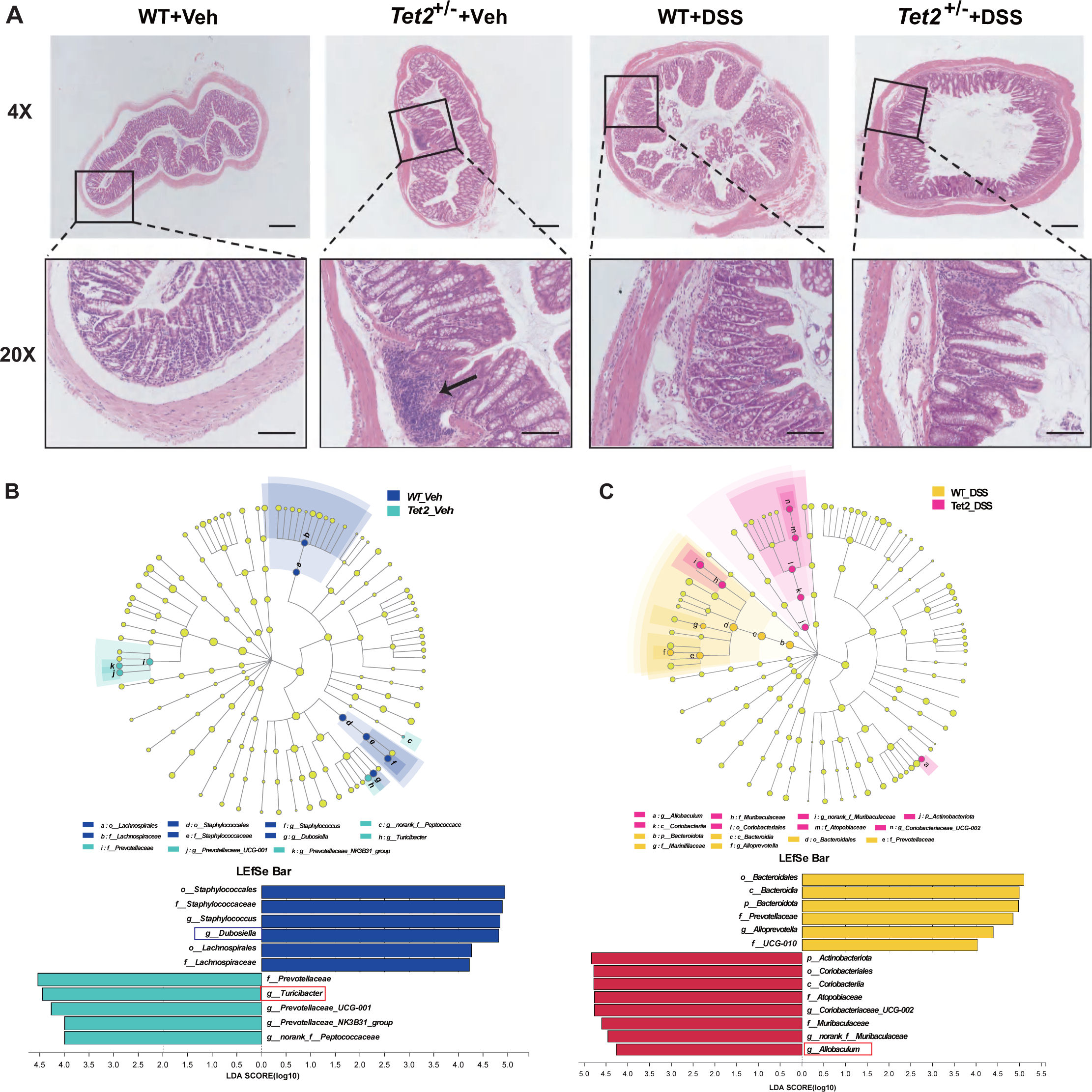

**Supplementary Figure 3.**
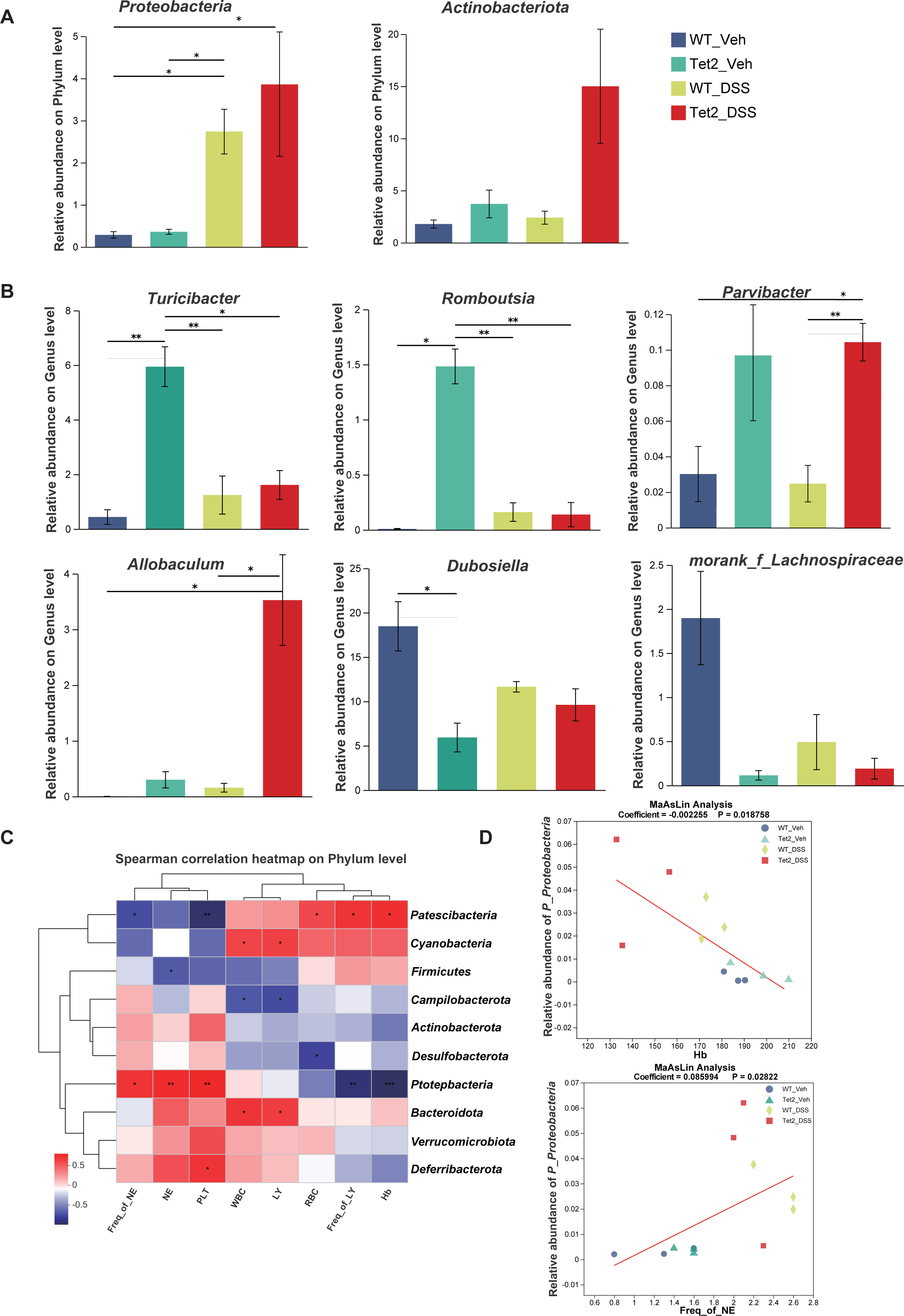

**Supplementary Figure 4.**
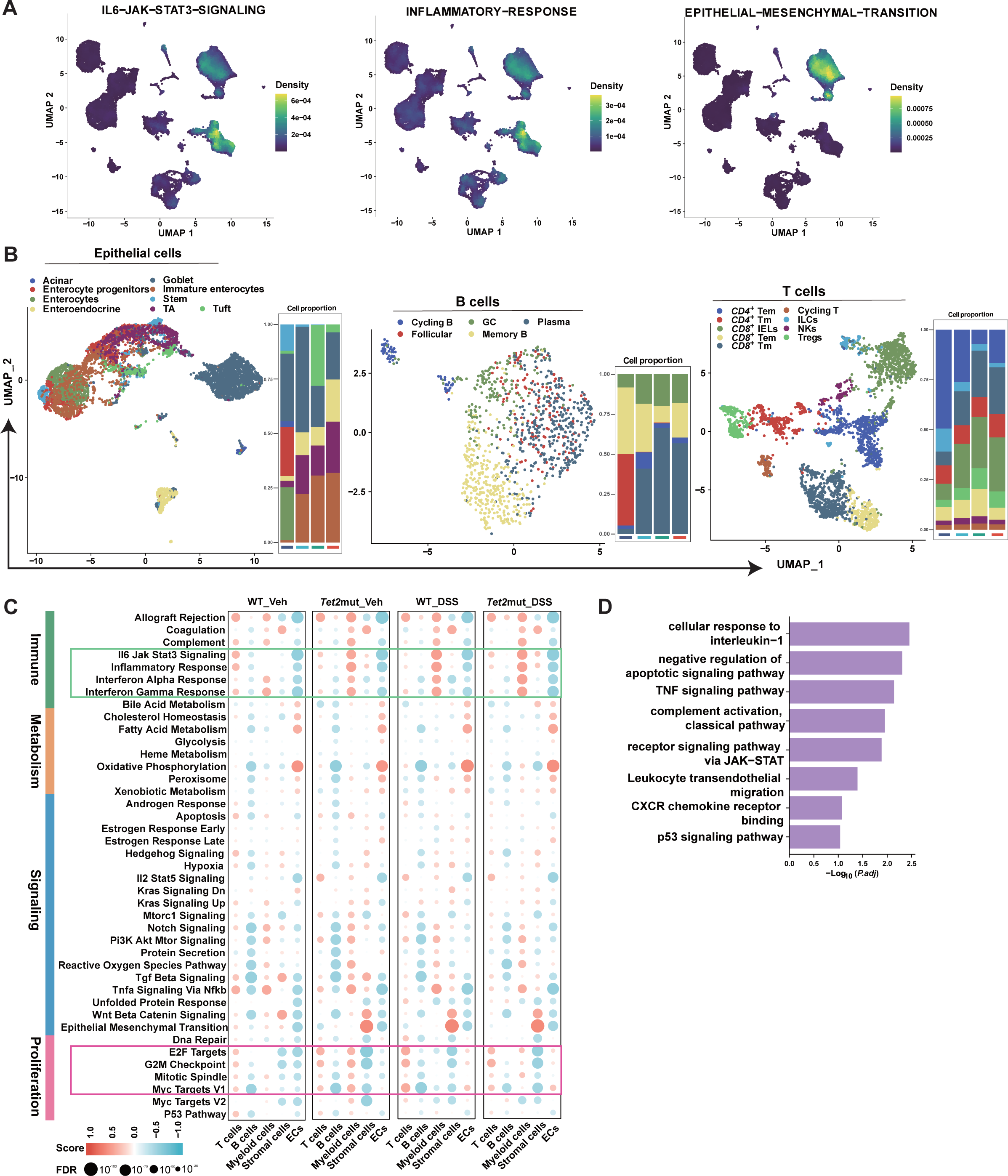

**Supplementary Figure 5.**
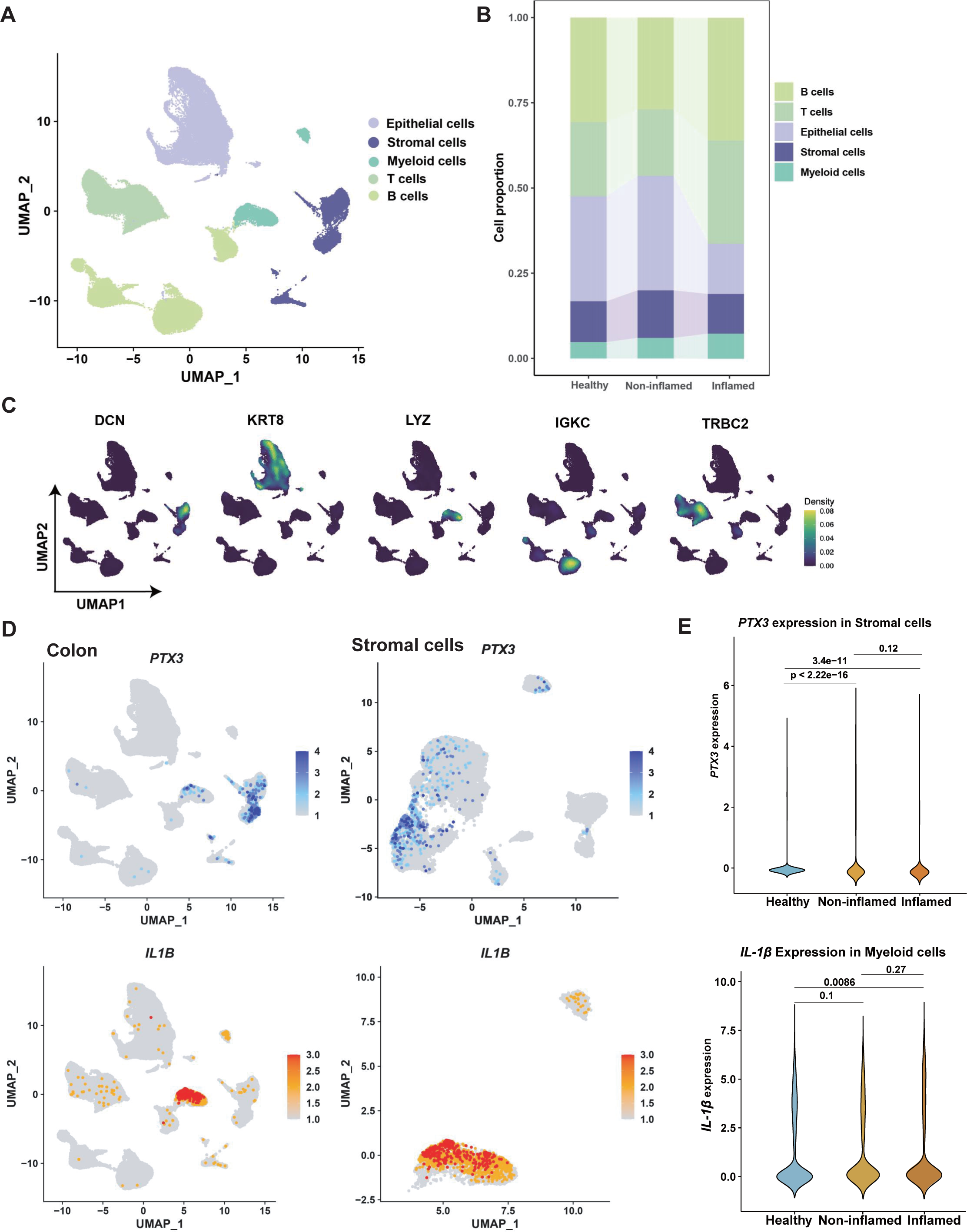

**Supplementary Figure 6.**
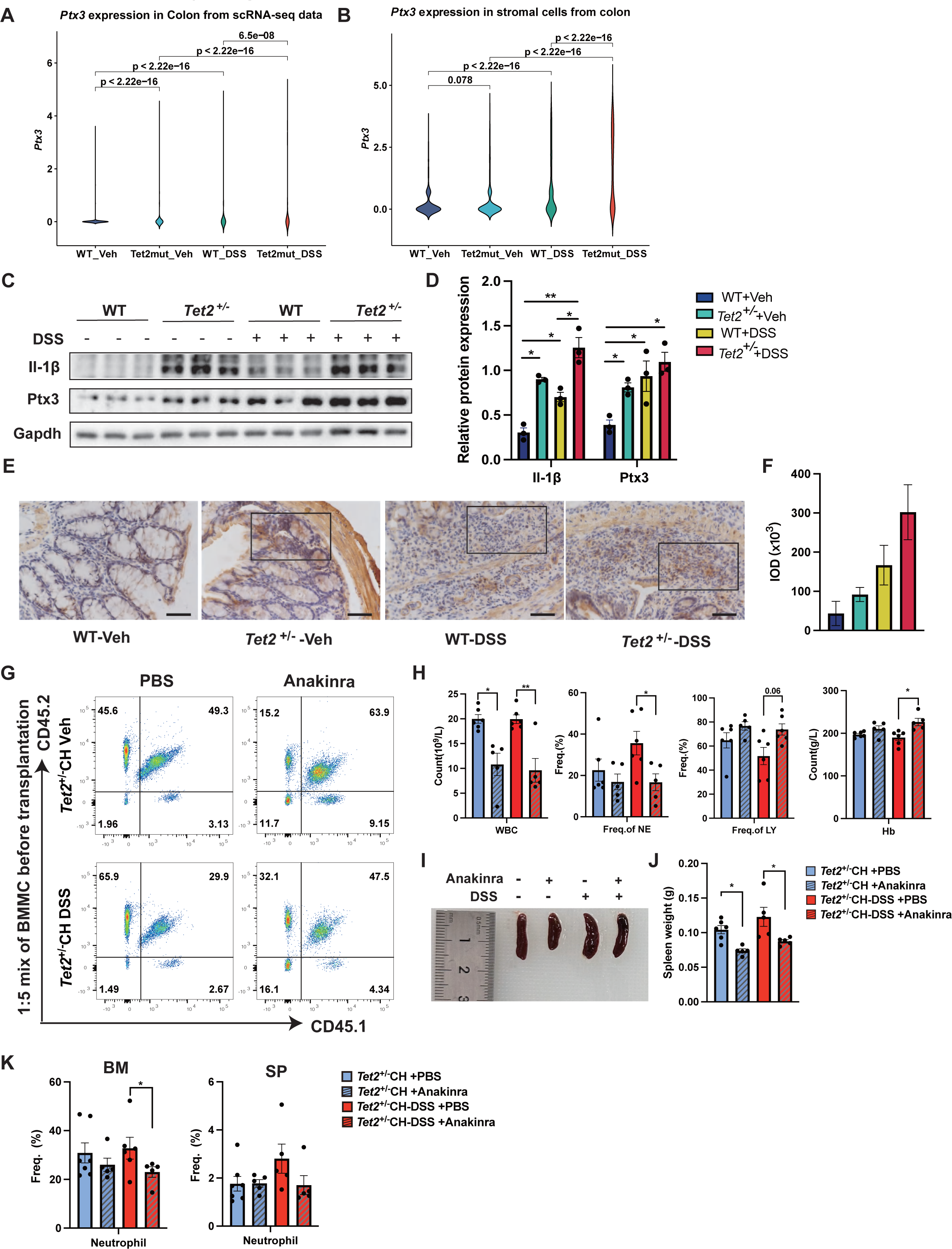

## Notes

### Competing Interest Statement

The authors have declared no competing interest.

### Summary of Updates

Figure 3 changed; Supplemental Figure 3 changed; Supplemental files updated.

## References

Abelson, S., Collord, G., Ng, S.W.K., Weissbrod, O., Mendelson Cohen, N., Niemeyer, E., Barda, N., Zuzarte, P.C., Heisler, L., Sundaravadanam, Y., et al. (2018). Prediction of acute myeloid leukaemia risk in healthy individuals. Nature 559, 400–404.

Agrawal, M., Niroula, A., Cunin, P., McConkey, M., Shkolnik, V., Kim, P.G., Wong, W.J., Weeks, L.D., Lin, A.E., Miller, P.G., et al. (2022). TET2-mutant clonal hematopoiesis and risk of gout. Blood 140, 1094–1103.

Bonavita, E., Gentile, S., Rubino, M., Maina, V., Papait, R., Kunderfranco, P., Greco, C., Feruglio, F., Molgora, M., Laface, I., et al. (2015). PTX3 is an extrinsic oncosuppressor regulating complement-dependent inflammation in cancer. Cell 160, 700–714.

Browaeys, R., Saelens, W., and Saeys, Y. (2020). NicheNet: modeling intercellular communication by linking ligands to target genes. Nat Methods 17, 159–162.

Burns, S.S., Kumar, R., Pasupuleti, S.K., So, K., Zhang, C., and Kapur, R. (2022). Il-1r1 drives leukemogenesis induced by Tet2 loss. Leukemia 36, 2531–2534.

Cai, Z., Aguilera, F., Ramdas, B., Daulatabad, S.V., Srivastava, R., Kotzin, J.J., Carroll, M., Wertheim, G., Williams, A., Janga, S.C., et al. (2020a). Targeting Bim via a lncRNA Morrbid Regulates the Survival of Preleukemic and Leukemic Cells. Cell Rep 31, 107816.

Cai, Z., Kotzin, J.J., Ramdas, B., Chen, S., Nelanuthala, S., Palam, L.R., Pandey, R., Mali, R.S., Liu, Y., Kelley, M.R., et al. (2018). Inhibition of Inflammatory Signaling in Tet2 Mutant Preleukemic Cells Mitigates Stress-Induced Abnormalities and Clonal Hematopoiesis. Cell Stem Cell 23, 833–849.e835.

Cai, Z., Lu, X., Zhang, C., Nelanuthala, S., Aguilera, F., Hadley, A., Ramdas, B., Fang, F., Nephew, K., Kotzin, J.J., et al. (2021). Hyperglycemia cooperates with Tet2 heterozygosity to induce leukemia driven by proinflammatory cytokine-induced lncRNA Morrbid. J Clin Invest 131.

Cai, Z., Zhang, C., Kotzin, J.J., Williams, A., Henao-Mejia, J., and Kapur, R. (2020b). Role of lncRNA Morrbid in PTPN11(Shp2)E76K-driven juvenile myelomonocytic leukemia. Blood Adv 4, 3246–3251.

Caiado, F., Kovtonyuk, L.V., Gonullu, N.G., Fullin, J., Boettcher, S., and Manz, M.G. (2023). Aging drives Tet2+/- clonal hematopoiesis via IL-1 signaling. Blood 141, 886–903.

Caiado, F., Pietras, E.M., and Manz, M.G. (2021). Inflammation as a regulator of hematopoietic stem cell function in disease, aging, and clonal selection. J Exp Med 218.

Chu, S.H., Heiser, D., Li, L., Kaplan, I., Collector, M., Huso, D., Sharkis, S.J., Civin, C., and Small, D. (2012). FLT3-ITD knockin impairs hematopoietic stem cell quiescence/homeostasis, leading to myeloproliferative neoplasm. Cell Stem Cell 11, 346–358.

Cumbo, C., Tarantini, F., Zagaria, A., Anelli, L., Minervini, C.F., Coccaro, N., Tota, G., Impera, L., Parciante, E., Conserva, M.R., et al. (2022). Clonal Hematopoiesis at the Crossroads of Inflammatory Bowel Diseases and Hematological Malignancies: A Biological Link? Front Oncol 12, 873896.

Delhommeau, F., Dupont, S., Della Valle, V., James, C., Trannoy, S., Massé, A., Kosmider, O., Le Couedic, J.P., Robert, F., Alberdi, A., et al. (2009). Mutation in TET2 in myeloid cancers. N Engl J Med 360, 2289–2301.

Deshmukh, H.S., Liu, Y., Menkiti, O.R., Mei, J., Dai, N., O’Leary, C.E., Oliver, P.M., Kolls, J.K., Weiser, J.N., and Worthen, G.S. (2014). The microbiota regulates neutrophil homeostasis and host resistance to Escherichia coli K1 sepsis in neonatal mice. Nat Med 20, 524–530.

Fabre, M.A., de Almeida, J.G., Fiorillo, E., Mitchell, E., Damaskou, A., Rak, J., Orrù, V., Marongiu, M., Chapman, M.S., Vijayabaskar, M.S., et al. (2022). The longitudinal dynamics and natural history of clonal haematopoiesis. Nature 606, 335–342.

Feng, Y., Yuan, Q., Newsome, R.C., Robinson, T., Bowman, R.L., Zuniga, A.N., Hall, K.N., Bernsten, C.M., Shabashvili, D.E., Krajcik, K.I., et al. (2023). Hematopoietic-specific heterozygous loss of Dnmt3a exacerbates colitis-associated colon cancer. Journal of Experimental Medicine 220.

Genovese, G., Kähler, A.K., Handsaker, R.E., Lindberg, J., Rose, S.A., Bakhoum, S.F., Chambert, K., Mick, E., Neale, B.M., Fromer, M., et al. (2014). Clonal hematopoiesis and blood-cancer risk inferred from blood DNA sequence. N Engl J Med 371, 2477–2487.

He, H., Wang, Z., Yu, H., Zhang, G., Wen, Y., and Cai, Z. (2022). Prioritizing risk genes as novel stratification biomarkers for acute monocytic leukemia by integrative analysis. Discov Oncol 13, 55.

Heyde, A., Rohde, D., McAlpine, C.S., Zhang, S., Hoyer, F.F., Gerold, J.M., Cheek, D., Iwamoto, Y., Schloss, M.J., Vandoorne, K., et al. (2021). Increased stem cell proliferation in atherosclerosis accelerates clonal hematopoiesis. Cell 184, 1348–1361.e1322.

Hsu, J.I., Dayaram, T., Tovy, A., De Braekeleer, E., Jeong, M., Wang, F., Zhang, J., Heffernan, T.P., Gera, S., Kovacs, J.J., et al. (2018). PPM1D Mutations Drive Clonal Hematopoiesis in Response to Cytotoxic Chemotherapy. Cell Stem Cell 23, 700–713.e706.

Jaiswal, S., and Ebert, B.L. (2019). Clonal hematopoiesis in human aging and disease. Science 366.

Jaiswal, S., and Libby, P. (2020). Clonal haematopoiesis: connecting ageing and inflammation in cardiovascular disease. Nat Rev Cardiol 17, 137–144.

Jaiswal, S., Natarajan, P., Silver, A.J., Gibson, C.J., Bick, A.G., Shvartz, E., McConkey, M., Gupta, N., Gabriel, S., Ardissino, D., et al. (2017). Clonal Hematopoiesis and Risk of Atherosclerotic Cardiovascular Disease. N Engl J Med 377, 111–121.

Khan, N., Patel, D., Trivedi, C., Kavani, H., Pernes, T., Medvedeva, E., Lewis, J., Xie, D., and Yang, Y.X. (2021). Incidence of Acute Myeloid Leukemia and Myelodysplastic Syndrome in Patients With Inflammatory Bowel Disease and the Impact of Thiopurines on Their Risk. Am J Gastroenterol 116, 741–747.

Khoury, J.D., Solary, E., Abla, O., Akkari, Y., Alaggio, R., Apperley, J.F., Bejar, R., Berti, E., Busque, L., Chan, J.K.C., et al. (2022). The 5th edition of the World Health Organization Classification of Haematolymphoid Tumours: Myeloid and Histiocytic/Dendritic Neoplasms. Leukemia 36, 1703–1719.

Li, Z., Cai, X., Cai, C.-L., Wang, J., Zhang, W., Petersen, B.E., Yang, F.-C., and Xu, M. (2011). Deletion of Tet2 in mice leads to dysregulated hematopoietic stem cells and subsequent development of myeloid malignancies. Blood 118, 4509–4518.

Libby, P., Ridker, P.M., and Maseri, A. (2002). Inflammation and atherosclerosis. Circulation 105, 1135–1143.

Meisel, M., Hinterleitner, R., Pacis, A., Chen, L., Earley, Z.M., Mayassi, T., Pierre, J.F., Ernest, J.D., Galipeau, H.J., Thuille, N., et al. (2018). Microbial signals drive pre-leukaemic myeloproliferation in a Tet2-deficient host. Nature 557, 580–584.

Miller, P.G., Qiao, D., Rojas-Quintero, J., Honigberg, M.C., Sperling, A.S., Gibson, C.J., Bick, A.G., Niroula, A., McConkey, M.E., Sandoval, B., et al. (2022). Association of clonal hematopoiesis with chronic obstructive pulmonary disease. Blood 139, 357–368.

Moran-Crusio, K., Reavie, L., Shih, A., Abdel-Wahab, O., Ndiaye-Lobry, D., Lobry, C., Figueroa, M.E., Vasanthakumar, A., Patel, J., Zhao, X., et al. (2011). Tet2 loss leads to increased hematopoietic stem cell self-renewal and myeloid transformation. Cancer Cell 20, 11–24.

Nazha, A., Komrokji, R., Meggendorfer, M., Jia, X., Radakovich, N., Shreve, J., Hilton, C.B., Nagata, Y., Hamilton, B.K., Mukherjee, S., et al. (2021). Personalized Prediction Model to Risk Stratify Patients With Myelodysplastic Syndromes. J Clin Oncol 39, 3737–3746.

Rasmussen, K.D., Jia, G., Johansen, J.V., Pedersen, M.T., Rapin, N., Bagger, F.O., Porse, B.T., Bernard, O.A., Christensen, J., and Helin, K. (2015). Loss of TET2 in hematopoietic cells leads to DNA hypermethylation of active enhancers and induction of leukemogenesis. Genes Dev 29, 910–922.

Smillie, C.S., Biton, M., Ordovas-Montanes, J., Sullivan, K.M., Burgin, G., Graham, D.B., Herbst, R.H., Rogel, N., Slyper, M., Waldman, J., et al. (2019). Intra- and Inter-cellular Rewiring of the Human Colon during Ulcerative Colitis. Cell 178, 714–730.e722.

Steensma, D.P., Bejar, R., Jaiswal, S., Lindsley, R.C., Sekeres, M.A., Hasserjian, R.P., and Ebert, B.L. (2015). Clonal hematopoiesis of indeterminate potential and its distinction from myelodysplastic syndromes. Blood 126, 9–16.

Taur, Y., Xavier, J.B., Lipuma, L., Ubeda, C., Goldberg, J., Gobourne, A., Lee, Y.J., Dubin, K.A., Socci, N.D., Viale, A., et al. (2012). Intestinal domination and the risk of bacteremia in patients undergoing allogeneic hematopoietic stem cell transplantation. Clin Infect Dis 55, 905–914.

Weeks, L.D., Niroula, A., Neuberg, D.S., Wong, W.J., Lindsley, R.C., Luskin, M.R., Berliner, N., Stone, R.M., DeAngelo, D.J., Soiffer, R.J., et al. (2022). Prediction of Risk for Myeloid Malignancy in Clonal Hematopoiesis. Blood 140, 2229–2231.

Weinstock, J.S., Gopakumar, J., Burugula, B.B., Uddin, M.M., Jahn, N., Belk, J.A., Bouzid, H., Daniel, B., Miao, Z., Ly, N., et al. (2023). Aberrant activation of TCL1A promotes stem cell expansion in clonal haematopoiesis. Nature 616, 755–763.

Wong, T.N., Ramsingh, G., Young, A.L., Miller, C.A., Touma, W., Welch, J.S., Lamprecht, T.L., Shen, D., Hundal, J., Fulton, R.S., et al. (2015). Role of TP53 mutations in the origin and evolution of therapy-related acute myeloid leukaemia. Nature 518, 552–555.

Yan, H., Baldridge, M.T., and King, K.Y. (2018). Hematopoiesis and the bacterial microbiome. Blood 132, 559–564.

Zeng, H., He, H., Guo, L., Li, J., Lee, M., Han, W., Guzman, A.G., Zang, S., Zhou, Y., Zhang, X., et al. (2019). Antibiotic treatment ameliorates Ten-eleven translocation 2 (TET2) loss-of-function associated hematological malignancies. Cancer Lett 467, 1–8.

Zeng, X., Li, X., Li, X., Wei, C., Shi, C., Hu, K., Kong, D., Luo, Q., Xu, Y., Shan, W., et al. (2023). Fecal microbiota transplantation from young mice rejuvenates aged hematopoietic stem cells by suppressing inflammation. Blood 141, 1691–1707.

Zhang, C.R.C., Nix, D., Gregory, M., Ciorba, M.A., Ostrander, E.L., Newberry, R.D., Spencer, D.H., and Challen, G.A. (2019). Inflammatory cytokines promote clonal hematopoiesis with specific mutations in ulcerative colitis patients. Exp Hematol 80, 36–41.e33.

Zhang, Q., Zhao, K., Shen, Q., Han, Y., Gu, Y., Li, X., Zhao, D., Liu, Y., Wang, C., Zhang, X., et al. (2015). Tet2 is required to resolve inflammation by recruiting Hdac2 to specifically repress IL-6. Nature 525, 389–393.

