## Supplemental Figure Legends for "Cooperative progression of colitis and leukemia modulated by clonal hematopoiesis via PTX3/IL-1β pro-inflammatory signaling"

**Supplemental Figure 1. Assessing impacts of various treatments on TedCH trajectory, related to Figure 1.**

(A-C) Representative flow profiles of PB from the chimeric mice treated with STZ (A), or 5-FU (B), or additional irradiation (C) as indicated.

Data are shown as means ± SEMs. Number of biological repeats (animals): n = 3~7. Experiments of DSS treatment on primary mice were repeated three times. *, p < 0.05; **, p < 0.01; ***, p < 0.001; ****, p < 0.0001.

**Supplemental Figure 2. *Tet2* deficiency exacerbated DSS-induced chronic colitis, related to Figure 2.**

(A) Images of hematoxylin and eosin (H&E) staining of spleen in each group after DSS treatment. (Scale bar: 4x, 500μm; 20x, 100μm).

(B-C) Cladogram generated from linear discriminant analysis effect size (LEfSe) and. the LDA score.

**Supplemental Figure3 Tet2 deficiency leads to abnormal intestinal barrier function, related to Figure 2.**

(A-B) The relative proportions of characteristic gut microbiota at phylum level and genus level.

(C) Heatmap of the Spearman correlation analysis between the gut microbiota and blood physiological values.

(D) MaAsLin analysis between the relative abundance of *proteobacteria* and hemoglobin or frequency of neutrophils.

Data are shown as means ± SEMs. Number of biological repeats (animals): n = 3~7. Experiments of DSS treatment on primary mice were repeated three times. *, p < 0.05; **, p < 0.01; ***, p < 0.001; ****, p < 0.0001.

**Supplemental Figure 4. Characterization of colon cells in *Tet2^+/-^* mice after chronic infection, related to Figure 4.**

(A) UMAP visualization of enrichment scores for specific gene sets.

(B) UMAP showing the composition of epithelial cells, B cells and T cells colored by cluster (left), and Bar plot showing the changes in the cell proportion of each group (right).

(C) Heatmap of dysregulated biological pathways in the 5 populations of colon tissues from 4 groups of samples.

(D) Representative KEGG pathways enrichment of the predicted target genes expressed in *Ptx3^+^* ﬁbroblasts.

**Supplemental Figure 5. Aberrant PTX3/IL-1β signaling is indicated in human colitis through analysis of clinical samples, related to Figure 5.**

(A) UMAP plot showing 5 main populations in colon tissues. A total of 90454 high-quality cells with an average of about 3000 genes per cell were included in the UMAP plot. After quality control during the dataset analysis, the group of *Health* has 55897 cells; *Non-inflamed* has 20188 cells; *Inflamed* has 14369 cells.

(B) Stacking bar plot showing the portion of the 5 main annotated populations in each colon sample.

(C) Expression of representative annotation markers for the 5 main cell populations in the UMAP plot of colons. *DCN* for stromal cells; *KRT*8 for epithelial cells; *LYZ* for myeloid cells; *IGKC* for B cells; and *TRBC2* for T cells.

(D) Expression of *PTX3* and *IL-1β* in the colon tissues (left panel) or in stromal cells (up-right panel) or in myeloid cells (bottom-left panel), respectively.

(E) Violin plots showing the expression of *PTX3* in the stromal cells (up panel) or in stromal cells (bottom panel).

**Supplemental Figure 6. The expression of Ptx3 and Il-1β was increased in the colon of *Tet2^+/-^* mice with chronic infection, related to Figure 6.**

(A-B) Expression of *Ptx3* in total colon tissues (A) and stromal cells (B) from the colon scRNA-seq dataset.

(C-D) The proteins level of Il-1β and Ptx3 were measured in colon tissues and quantified.

(E-F) Representative immunohistochemical (IHC) staining of Ptx3 in colon (E) and semi-quantitative analysis of immunohistochemistry staining of Ptx3 based on integrated optical density (IOD) (F). scale bar is 100 μm.

(G) Representative flow profiles of PB from the chimeric mice treated with Anakinra or PBS.

(H) Hematological parameters of PB were monitored at the end-point of the Anakinra treatment.

(I-J) Photography (I) and quantification of spleens (J) for the TedCH chimeric mice treated with DSS and/or Anakinra.

(K) Quantification of mature cells including neutrophils in bone marrow and spleen. BM, bone marrow; SP, spleen; PB, peripheral blood.

Data are shown as means ± SEMs. Number of biological repeats (animals): n = 5~7. Experiments of Anakinra treatment were repeated twice. *, p < 0.05; **, p < 0.01.
